# Dissociable changes in temporal precision and causal expectations following perceptual training

**DOI:** 10.64898/2026.09.06.749727

**Authors:** Haocheng Zhu, Shizhong Cai, Jingci Lang, Xiaole Liu, Aijun Wang

## Abstract

A common behavioral gain does not necessarily reflect a common learning mechanism. Perceptual learning may sharpen sensory evidence without updating higher-level expectations about its causes. Across three experiments, we examined audiovisual binding and its plasticity in children with ADHD and typically developing children. Children with ADHD were more susceptible to the sound-induced flash illusion (Experiment 1) and showed markedly flatter temporal tuning of it (Experiment 2), yet conventional temporal binding window (TBW) estimates did not distinguish the groups—behavior alone could not localize the difference. Following six days of simultaneity-judgment training, both groups became comparably less susceptible to the illusion. A temporally explicit two-dimensional Bayesian causal inference model, however, revealed sharply different latent changes. Typically developing children reduced their causal prior expectation that auditory and visual signals shared a common cause. Children with ADHD showed no corresponding prior recalibration but instead exhibited a marked reduction in temporal uncertainty, indicating more precise perceptual evidence that did not propagate to the slower, more abstract estimate of environmental causal structure. Thus, perceptual learning converged at the behavioral level while diverging computationally. These findings argue against unitary accounts of multisensory plasticity and show how the same perceptual endpoint can conceal adaptation at different levels of the inferential hierarchy.

## 1. Introduction

Learning to perceive better can mean two quite different things. An observer may come to register the incoming evidence more precisely, or may revise their beliefs about the world that generated it. Perception depends on both, being an inference in which current evidence is weighed against expectations about how the world tends to be arranged (Körding et al., 2007; Shams & Beierholm, 2010, 2022; Noppeney, 2021), and experience can act on either. These forms of plasticity need not unfold together. More precise sensory evidence can change how an event is perceived without necessarily revising the expectations brought to subsequent events. One possibility is that expectations about environmental causal structure are updated over a longer timescale than the precision of sensory evidence, allowing perceptual improvement to precede a measurable change in those expectations. Learning could therefore improve the information available to the observer while leaving a higher-level belief relatively unchanged. This raises a question about the coordination of learning across levels of inference: does improved sensory evidence lead to revised causal expectations, or can the two remain dissociated over the same period of experience?

Multisensory perception is one of the few domains in which these ingredients have been formalized well to be estimated apart. Whether a sound and a flash are bound into a single event depends jointly on how precisely each signal is encoded, how precisely their relative timing is represented, how strongly the observer expects signals in different modalities to have arisen from a common cause, and the rule by which the resulting posterior is turned into a response (Körding et al., 2007; Wozny et al., 2010; Rohe & Noppeney, 2015; Rohe et al., 2019; Kayser & Shams, 2015; Zhu et al., 2024a). Behavioral modelling provides evidence that causal priors can remain stable across changes in sensory likelihoods (Beierholm et al., 2009). Crucially, sharper sensory evidence can alter the posterior probability of a common cause on a given trial even when the common-cause prior remains unchanged. The question here concerns whether experience also produces a lasting revision of that prior. Behavioral summary measures, however, do not uniquely distinguish changes in sensory precision from changes in causal expectations. The temporal binding window, the range of asynchronies over which signals still influence one another (Stevenson et al., 2012; Wallace & Stevenson, 2014; Stevenson et al., 2017), narrows whether the observer’s temporal evidence became sharper or the observer’s expectation of a common cause weakened; susceptibility to the sound-induced flash illusion falls under either change as well (Zhu et al., 2024a, 2026a). Neither measure alone establishes whether perceptual improvement is accompanied by a lasting revision of the observer’s causal expectations.

The distinction can be drawn when the components are modelled rather than summarized, and a number of studies have now done so. Modality-specific attention attenuates integration and recalibration specifically by lowering the prior expectation of a common source, leaving sensory precision intact (Badde et al., 2020). Behavioral simultaneity windows track temporal sensory uncertainty rather than a dedicated tolerance parameter (Cary et al., 2024). Causal inference gates recalibration itself (Hong et al., 2021; Li et al., 2025). Perceptual training has also been modelled as jointly increasing the precision of audiovisual timing estimates and reducing the common-cause prior (McGovern et al., 2016). Group differences submit to the same treatment: atypical multisensory perception in autism has been localized to altered causal priors rather than uniformly degraded precision (Noel et al., 2022, 2023; Baum et al., 2015; Foss-Feig et al., 2010; Stevenson et al., 2014), while causal inference appears largely preserved in medicated post-acute psychosis (Rohe et al., 2024). These findings establish that sensory precision and causal expectations can each contribute to perceptual change. Their co-occurrence, however, does not establish that an improvement in sensory evidence necessarily leads to an update of causal expectations. A further question is whether these forms of plasticity remain coordinated across learners. If causal expectations update more slowly, improved sensory precision may be detectable during a period in which the common-cause prior shows little measurable change. Separately estimating the two components therefore provides a way to test for a dissociation between perceptual improvement and the revision of causal expectations.

Perceptual training provides a setting in which to ask whether improvements in sensory evidence are accompanied by changes in causal expectations over a defined period of experience. A few sessions of feedback on audiovisual simultaneity are enough to change how adults combine sight and sound (Powers et al., 2009; Setti et al., 2014), with corresponding modulation of auditory and multisensory cortical responses (Powers et al., 2012). Outside healthy adults, however, the picture fragments. A randomized controlled trial in autistic youth found benefits confined to those with higher cognitive ability, with paradoxical widening in those with co-occurring impairments (Feldman et al., 2023). In older adults, the same protocol narrowed the window but left largely untouched the crossmodal interaction patterns that plausibly index prior expectations (McGovern et al., 2022). Audiovisual temporal sensitivity and cumulative recalibration also mature on different schedules (Hillock-Dunn & Wallace, 2012; Rohlf et al., 2020; Ampollini et al., 2024). These findings motivate separating changes in temporal sensitivity from changes in causal expectations when evaluating training. A behavioral gain could reflect changes in either or both components. If the components adapt over different timescales, an assessment after a fixed period of training may reveal improved sensory precision before a reliable change in the causal prior becomes apparent.

The sound-induced flash illusion provides a suitable setting in which to make the test. A single flash accompanied by two beeps is frequently reported as two flashes (Shams et al., 2000, 2002), and because the illusion is graded by audiovisual asynchrony it is sensitive to the numerosity and the timing dimensions of the inference simultaneously (Hirst et al., 2020). It is also among the few multisensory phenomena with a worked-out generative account, from the original optimal-percept treatment (Shams et al., 2005) through explicitly temporal two-dimensional formulations (Zhu et al., 2026b) to competing non-Bayesian time-window models (Colonius & Diederich, 2025). Crucially for present purposes, a model of this kind returns separate estimates of numerosity precision, temporal precision and the common-cause prior from the same behavioral data, so that a change in illusion susceptibility can be assigned to a component rather than merely recorded (Shams et al., 2002; Wang et al., 2025; Zhu et al., 2026a).

Children with ADHD provide a theoretically informative comparison for examining whether changes in temporal precision and causal expectations can dissociate. Although ADHD has traditionally been characterized through executive dysfunction (Barkley, 1997; Willcutt et al., 2005; Faraone et al., 2015), sensory atypicality is pervasive and meta-analytically confirmed (Ghanizadeh, 2011; Shimizu et al., 2014; Lane & Reynolds, 2019; Jurek et al., 2025), with consequences for behaviour in cluttered and distracting environments (Yochman et al., 2004; Koziol & Budding, 2012). More specifically, millisecond-range timing is both less accurate and more variable across a range of tasks (Smith et al., 2002; Toplak et al., 2006; Marusich & Gilden, 2014), a pattern borne out by two meta-analyses, though with magnitude and direction depending on the temporal operation measured (Zheng et al., 2022; Marx et al., 2022); ADHD-like traits in adults have similarly been linked to altered audiovisual temporal integration (Panagiotidi et al., 2017). This yields a directional expectation about one ingredient of the inference, elevated temporal uncertainty at intake, and no comparable expectation about the other. That asymmetry is what makes the comparison diagnostic rather than merely descriptive. It also cannot be reduced to a general learning deficit, where rapid visual statistical learning and the use of low-level perceptual priors are intact in adults with ADHD (Richards et al., 2020), and reward learning in ADHD has been associated with excessive rather than insufficient switching (Aster et al., 2024). Whatever inflexibility means in this context (Dajani & Uddin, 2015), it has not been located within perceptual causal inference, and locating it is exactly the kind of question a decomposed model can make precise.

Two accounts of what should happen follow directly. On a fixed-locus account, a training regimen acts on a characteristic component of the computation, and both groups should show the same parameter change irrespective of where they began. On an opportunistic account, learning is expressed at whichever level currently limits performance, so children entering with elevated temporal uncertainty should show a reduction in temporal uncertainty, whereas children whose temporal evidence is already relatively precise should instead show recalibration of the common-cause prior. The two accounts make the same prediction for illusion susceptibility. They differ only in the latent quantity that moves, and are therefore separable only by a model that estimates the components individually.

Across three experiments, we examined whether comparable behavioral improvements could be accompanied by a dissociation between changes in temporal uncertainty and changes in the common-cause prior. Experiment 1 compared susceptibility to the fission and fusion illusions between children with ADHD and typically developing children. Experiment 2 varied audiovisual asynchrony to characterize the temporal profile of the fission illusion and to derive conventional binding window estimates. Experiment 3 gave both groups six sessions of adaptive simultaneity-judgement training, measured the illusion, simultaneity judgements and temporal order judgements before and after, and fitted a two-dimensional Bayesian causal inference model that estimates visual numerosity uncertainty, audiovisual temporal uncertainty and the common-cause prior separately, alongside the decision rule.

The contribution is not that priors and sensory precision are separable in principle; that much is established (Beierholm et al., 2009; Badde et al., 2020; Zhu et al., 2024b). It is that a single training protocol moved different components in different learners while producing the same behavioral outcome, which makes the locus of perceptual learning a property of the learner rather than of the regimen, and places a corresponding limit on what any behavioral endpoint can be taken to show. This dissociation suggests that behavioral improvement need not be accompanied by coordinated plasticity across levels of perceptual inference. It is consistent with the possibility that causal expectations update more slowly or less readily than sensory precision, leaving a higher-level expectation relatively stable despite improved perceptual evidence.

## 2. Methods

### 2.1. Ethics and shared eligibility criteria

All parents or legal guardians gave written informed consent in compliance with the Declaration of Helsinki, and children provided assent in a developmentally appropriate manner. This study was approved by the academic committee of the Department of Psychology at the host institution (approval number withheld for anonymous review). Children in the attention-deficit/hyperactivity disorder (ADHD) groups were recruited recruited through the child and adolescent healthcare department of a regional children’s hospital.

Eligibility criteria were shared across experiments. Children assigned to the ADHD group had received a clinical diagnosis of ADHD from a hospital clinician according to the *Diagnostic and Statistical Manual of Mental Disorders*, fifth edition (DSM-5; American Psychiatric Association, 2013). Additional inclusion criteria were right-handedness, normal hearing, normal or corrected-to-normal vision, and nonverbal reasoning above the 25th percentile on the Combined Raven’s Test. Children were excluded if they had a history of another neurological or psychiatric disorder. The ADHD samples were medication-naïve at enrolment. TD children met the same sensory, handedness, cognitive, and neurological criteria, had no ADHD diagnosis, and had no marked learning or behavioural problems according to parent and teacher interviews.

Unless specified otherwise, visual stimuli were presented using Presentation software (Neurobehavioral Systems, Berkeley, CA) against a black background on a Dell monitor (1024 × 768 pixels; 60-Hz refresh rate). Auditory stimuli were delivered binaurally through Audio-Technica ATH-WS99 headphones. The visual event was a white disc subtending approximately 2° of visual angle, presented 5° below a central fixation point for 17 ms. The auditory event was a 3.5-kHz, 70-dB pure tone with a duration of 7 ms. Children were instructed to maintain fixation throughout each trial. Trials were presented in randomized order, and no response deadline was imposed.

### 2.2. Experiment 1: susceptibility to the sound-induced flash illusion

#### 2.2.1. Participants

An a priori power analysis was conducted in G*Power 3.1.9.2 (Faul et al., 2007). Assuming a medium effect (*f* = 0.25), α = 0.01, and 80% power, the minimum total sample was 36 children. 56 children completed Experiment 1: 28 children with ADHD (16 boys and 12 girls; mean age = 8.9 years, SD = 1.80) and 28 TD children (13 boys and 15 girls; mean age = 9.3 years, SD = 1.54). No participant was excluded from the final Experiment 1 sample.

#### 2.2.2. Design and procedure

Experiment 1 used the visual-report version of the sound-induced flash illusion (SIFI; Shams et al., 2002). On every trial, children reported whether they perceived one or two flashes while ignoring any accompanying tones. Six stimulus conditions were presented. Two unimodal conditions contained one flash (F1) or two flashes (F2). Two audiovisually congruent conditions paired one flash with one tone (F1B1) or two flashes with two tones (F2B2). The two incongruent conditions paired one flash with two tones (F1B2; fission) or two flashes with one tone (F2B1; fusion). In multisensory conditions, the first flash and tone were temporally aligned. Each condition was presented 20 times, giving 120 trials in total. Trial order was randomized separately for each participant. Responses indicated whether one or two flashes had been perceived; failures to respond were coded as omissions.

The primary dependent variables were the fission-illusion rate, defined as the proportion of valid F1B2 trials on which two flashes were reported, and the fusion-illusion rate, defined as the proportion of valid F2B1 trials on which one flash was reported. Accuracy was additionally calculated for all six stimulus conditions.

#### 2.2.3. Behavioural analysis

Primary analyses were performed on valid responses; omission rates were calculated separately. Group differences in fission and fusion rates were evaluated with two-sided Welch independent-samples *t*-tests. For each comparison, the ADHD-minus-TD mean difference, its 95% confidence interval, and Cohen’s *d* were calculated. Two-sided Mann–Whitney tests were used as distribution-robust sensitivity analyses. To test whether any group difference was specific to illusion type, each participant’s fission-minus-fusion contrast was calculated and the contrasts were compared between groups using a Welch *t*-test.

Signal-detection measures were used to distinguish illusion-consistent perceptual reports from a general tendency to select the same response. For fission, the hit rate was the probability of reporting two flashes in F1B2 trials and the false-alarm rate was the probability of reporting two flashes in matched F1B1 trials. For fusion, the hit rate was the probability of reporting one flash in F2B1 trials and the false-alarm rate was the probability of reporting one flash in matched F2B2 trials. Sensitivity and criterion were calculated as

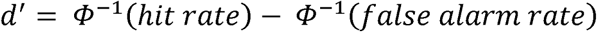

And

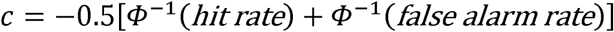

where Φ⁻¹ denotes the inverse cumulative standard-normal distribution. Extreme proportions were adjusted to 0.5/*n* or 1 − 0.5/*n*, where *n* was the number of valid trials in the corresponding condition (Macmillan and Kaplan, 1985; Stanislaw and Todorov, 1999). Group differences in *d*′ and *c* were tested with two-sided Welch *t*-tests, with Mann–Whitney tests reported as sensitivity checks. Condition-wise accuracy was compared between groups using Welch *t*-tests, with Holm adjustment across the six conditions. Missing-response rates were compared between groups. As an additional conservative analysis, fission and fusion rates were recomputed with all presented trials retained in the denominator, so that omissions could not inflate estimated illusion susceptibility.

### 2.3. Experiment 2: temporal modulation of the fission illusion

#### 2.3.1. Participants

The Experiment 2 sample-size calculation was performed in G*Power 3.1.9.2. Because temporal-window characterization involved within-participant comparisons across stimulus onset asynchronies (SOAs), the calculation was based on a paired comparison. A medium expected effect (*d*_z_ = 0.55), α = 0.05, and 80% power yielded a minimum of 28 children per group. Sixty children completed the experiment: 30 children with ADHD (21 boys and 9 girls; mean age = 9.5 years, SD = 1.90) and 30 TD children (21 boys and 9 girls; mean age = 10.2 years, SD = 1.30). The groups did not differ in age, *t*(58) = 1.66, *p* = 0.102, or sex distribution, χ²(1) = 0, *p* = 1.000. No participant was excluded from the final Experiment 2 sample.

#### 2.3.2. Design and procedure

The apparatus, basic stimuli, and visual numerosity report were the same as in Experiment 1. The principal manipulation varied the SOA in fission trials containing one flash and two tones (F1B2). One tone was synchronous with the flash, whereas the second tone occurred at one of ten non-zero SOAs: −300, −200, −150, −100, −50, +50, +100, +150, +200, or +300 ms. Negative values indicate that the asynchronous tone preceded the synchronous flash–tone pair (auditory-leading); positive values indicate that it followed the pair (visual-leading). Each SOA was presented 20 times, yielding 200 F1B2 trials.

The same ten SOAs were used in 200 congruent two-flash/two-tone trials (F2B2), in which the two audiovisual pairs were separated by the designated SOA. Four additional control conditions, F1, F1B1, F2, and F2B1, were each presented 20 times. The complete task therefore comprised 480 trials, randomly intermixed and divided into four blocks. Children could take a self-paced break between blocks. They pressed “F” to report one perceived flash and “J” to report two perceived flashes.

#### 2.3.3. Behavioural analysis

Omissions were excluded from the primary rate calculations. For each participant and SOA, the fission-illusion rate was the proportion of valid F1B2 trials on which two flashes were reported. The primary omnibus analysis was a 2 × 10 mixed analysis of variance with group (ADHD, TD) as a between-participant factor and SOA as a within-participant factor. Greenhouse–Geisser correction was applied to the SOA main effect and Group × SOA interaction. Significant or theoretically informative effects were followed with two-sided Welch *t*-tests at each SOA; the ten resulting *p*-values were Holm-adjusted.

Temporal modulation was quantified for each participant as the ordinary least-squares slope of fission-illusion rate against absolute SOA, expressed per 100 ms. More negative slopes indicate a steeper decline in illusion susceptibility as audiovisual separation increased; slopes nearer zero indicate flatter temporal tuning. We also calculated the mean illusion rate at near SOAs (|SOA| ≤ 100 ms), the mean rate at far SOAs (|SOA| ≥ 200 ms), and their difference. Group comparisons used two-sided Welch *t*-tests with 95% confidence intervals and Cohen’s *d*; Mann–Whitney tests provided sensitivity checks. As a model-free summary of temporal-window width, each participant’s SOA function was linearly interpolated at 1-ms resolution. TBW₈₀ was defined as the span between the first and last interpolated SOA at which the illusion rate was at least 80% of that participant’s observed peak rate. TBW₈₀ was treated as an exploratory descriptive measure because it depends on the finite SOA range and on sampling variability near the peak.

For comparison with conventional psychometric summaries, an exploratory four-parameter Gaussian was also fitted to each participant’s illusion-by-SOA function:

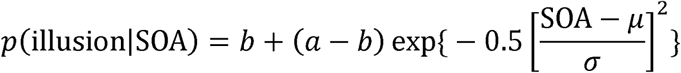

where *a* is the peak, *b* the baseline, μ the curve centre, and σ the Gaussian width. Fits were constrained to 0 ≤ *a*, *b* ≤ 1, 10 ≤ σ ≤ 500 ms, and −300 ≤ μ ≤ 300 ms. Fit quality was summarized by *R*². Gaussian parameters were not used to define the primary temporal-window result.

To compare the illusion with a participant-specific response baseline, the F1B2 rate at each SOA was contrasted with the F1B1 error rate using paired, two-sided tests; Holm correction was applied separately across the ten SOAs within each group. Error rates in the F1, F2, F1B1, and F2B2 control conditions were compared between groups with Welch *t*-tests and Holm adjustment across the four controls. Missing-response rates were also compared between groups.

### 2.4. Experiment 3: perceptual training and transfer across temporal tasks

#### 2.4.1. Participants

An a priori calculation in G*Power 3.1.9.7 was based on the within-participant pre–post training contrast. Assuming *d*_z_ = 0.55, α = 0.05, and 80% power, at least 18 completers were required per group. Sixty children enrolled; 10 withdrew before completing the protocol and were excluded. The final sample comprised 23 children with ADHD (18 boys and 5 girls; mean age = 9.0 years, SD = 1.23) and 27 TD children (17 boys and 10 girls; mean age = 9.1 years, SD = 1.13).

#### 2.4.2. Study design

Experiment 3 used a two-group pretest–training–post-test design. Both groups completed the same six-session adaptive audiovisual simultaneity-judgment (SJ) training; there was no untrained or active-control condition. Before the first training session and after the final session, children completed three assessment tasks: the SOA-resolved SIFI task, a non-adaptive SJ task, and a temporal-order judgment (TOJ) task. The pre- and post-test implementations were identical within each task.

#### 2.4.3. Pre- and post-training assessment tasks

The SIFI assessment was identical to Experiment 2 and comprised 480 trials. The present behavioural analysis focused on F1B2 trials at the ten non-zero SOAs. At each SOA, fission susceptibility was the proportion of valid trials on which two flashes were reported. The participant-level pre–post endpoint was the mean fission-illusion rate across these trials.

In the SJ assessment, one flash and one tone were presented at 13 SOAs: −300, −250, −200, −150, −100, −50, 0, +50, +100, +150, +200, +250, and +300 ms. Each SOA was presented ten times, giving 130 trials. Negative SOAs indicate auditory-leading trials and positive SOAs visual-leading trials. Children reported whether the events were simultaneous or asynchronous; zero-SOA trials were physically simultaneous. The outcome at each SOA was the proportion of valid trials judged simultaneous.

The TOJ assessment used the same non-zero SOAs as the SJ assessment (±50 to ±300 ms in 50-ms increments), with ten repetitions per SOA and 120 trials in total. Children reported whether the visual or auditory event occurred first. The primary behavioural summaries were overall response accuracy and the proportion of visual-first responses at each SOA.

The assessment tasks used brief central visual and auditory events with the same 3.5-kHz tone and disc geometry as the training task. Responses that were absent or outside the two valid response categories were treated as omissions and excluded from rate estimation.

#### 2.4.4. Adaptive simultaneity-judgment training

Training was administered once per day over six sessions; each session lasted approximately 30 min. The task was programmed in Presentation and run at a viewing distance of 60 cm on a Dell monitor (1024 × 768 pixels; 120-Hz refresh rate). The visual stimulus was a white disc subtending approximately 2.2° against a black background, and the auditory stimulus was a 3.5-kHz, 70-dB tone.

Each trial began with a central fixation cross for 500 ms, followed by a blank interval of 500 ms and then the audiovisual pair. Children judged whether the events were simultaneous (“M”) or asynchronous (“Z”), after which immediate correctness feedback was presented. Positive and negative SOAs were trained in separate adaptive staircases. The staircase began at an absolute SOA of 300 ms, as indicated by the recorded training files, and used 25-ms steps bounded between 25 and 300 ms. After three consecutive correct responses, the absolute SOA decreased by 25 ms; after one incorrect response, it increased by 25 ms. Thus, visual-leading and auditory-leading staircases independently approached each child’s discrimination threshold. Physically synchronous baseline trials (SOA = 0) were interleaved to discourage a uniform “asynchronous” response but did not determine staircase level. For descriptive analyses of learning across the six sessions, the daily threshold for each SOA direction was defined from the final reversal levels of the staircase.

#### 2.4.5. Behavioural analysis

All pre–post analyses preserved participant pairing. For SIFI, the primary endpoint was the participant’s mean fission-illusion rate. For SJ, endpoints included the overall simultaneous-response rate and an individual temporal-window estimate. For TOJ, endpoints included overall accuracy, the proportion of visual-first responses, and an individual uncertainty-window estimate. Within each group, pre–post differences were tested using two-sided paired *t*-tests and reported with 95% confidence intervals and standardized paired effect sizes (*d*_z_). Group × Time effects were tested by comparing participant-level change scores between the ADHD and TD groups with two-sided Welch *t*-tests. Holm-adjusted *p*-values were calculated within each endpoint and across the full family of behavioural endpoint tests.

Psychometric curves were estimated separately for each participant, group, time point, and task after first averaging binary responses within SOA. SIFI and SJ curves were fitted with a baseline-plus-amplitude Gaussian,

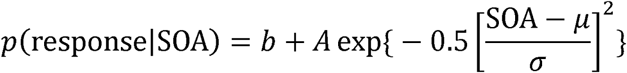

The fitted centre μ indexed the point of subjective simultaneity, and temporal-window width was defined as the full width at half maximum. Parameter bounds were 0 ≤ *b* ≤ 0.85, 0 ≤ *A* ≤ 1, −500 ≤ μ ≤ 500 ms, and 10 ≤ σ ≤ 1000 ms. SIFI curves were used to display the temporal profile of illusion susceptibility; the principal individual SIFI endpoint remained the observed fission rate.

TOJ curves were fitted with a cumulative Gaussian,

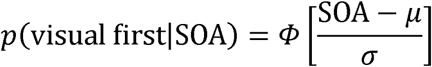

where μ is the point of subjective simultaneity and σ controls the slope. Bounds were −1500 ≤ μ ≤ 1500 ms and 10 ≤ σ ≤ 2500 ms. The TOJ uncertainty window was the distance between the 25% and 75% points of the fitted function. This estimate was retained for transparency even when it extended beyond the tested 600-ms SOA range; such extrapolated or boundary-limited estimates were interpreted cautiously.

Because each assessment contained relatively few repetitions per SOA, inference on group-level SJ and TOJ curve changes used a paired participant bootstrap rather than treating individual fits as error-free. For each task and group, participants were sampled with replacement while their pre–post pairing was preserved. The mean pre- and post-test curves were refitted on each of 3,000 bootstrap samples. Percentile 95% confidence intervals and two-sided bootstrap tail probabilities were obtained for pre–post changes in temporal-window width and curve center. Group differences in change were evaluated from the bootstrap distribution of the ADHD-minus-TD difference between the two group-specific changes.

### 2.5. Computational modelling

#### 2.5.1. Two-dimensional Bayesian causal inference model of the SIFI

We modelled trial-by-trial visual numerosity reports using a two-dimensional Bayesian causal inference (BCI) framework adapted from Zhu et al. (2026b). The model treats numerosity and relative timing as conditionally independent sources of evidence about a shared latent causal structure, *C*. Under the common-cause hypothesis (*C* = 1), the flashes and beeps are generated by one audiovisual event; under the separate-causes hypothesis (*C* = 2), the visual and auditory streams arise independently. The prior probability of a common cause, *p_common_* = *P*(*C* = 1), quantifies the observer’s expectation that the two streams belong together before the current sensory evidence is observed.

For each simulated trial, the physical numbers of flashes and beeps (*S_V_* and *S_A_*) generated noisy internal numerosity measurements (*x_V_* and *x_A_*). Visual and auditory onsets were represented relative to the visual event. Unlike the formulation with separate auditory and visual temporal-noise parameters, our identifiable parameterization used a single temporal uncertainty, σ*_t_*, for both modalities. An additional latency parameter, δ, shifted the auditory measurement and therefore allowed the center of the temporal-evidence function to depart from physical synchrony:

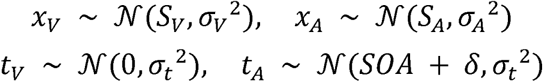

Numerosity sources followed a Gaussian prior, 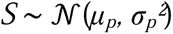, and temporal sources followed φ 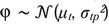. For the primary model, μ*_t_* was fixed at 0 ms and σ*_tp_* at 300 ms. For each dimension, evidence under *C* = 1 was obtained by integrating the two modality-specific likelihoods over one latent source; evidence under *C* = 2 was the product of two corresponding single-modality integrals. Following Zhu et al. (2026b), numerosity and temporal measurements were assumed to be conditionally independent given the causal structure. Bayes’ rule therefore gives

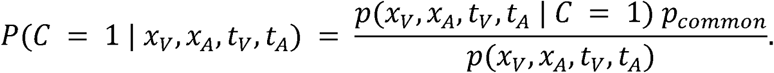

To make the contribution of the two dimensions explicit, let *L_N_*⁽□⁾ = *p*(*x_V_*, *x_A_* | *C* = *c*) denote the numerosity likelihood and *L_T_*⁽□⁾ = *p*(*t_V_*, *t_A_*| *C* = *c*) the temporal likelihood. Expanding the marginal evidence in the denominator gives the probability-space expression used by the multidimensional model:

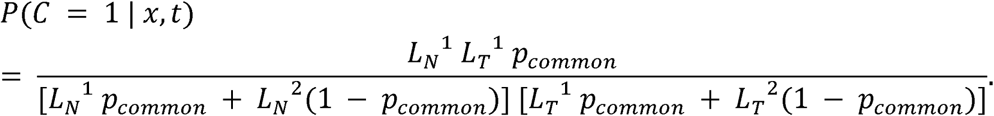

This posterior is the model’s trial-wise probability that the flashes and beeps arose from a common event. Values near 1 favor integration; values near 0 favor segregation. In the code, the same posterior was evaluated from log likelihood ratios using numerically stable logistic transformations, which is an algebraically equivalent implementation of the probability-space equations above.

Under *C* = 1, the fused numerosity estimate combined *x_V_*, *x_A_*, and the numerosity prior in proportion to their precisions. Under *C* = 2, the visual estimate combined only *x_V_*with the numerosity prior. Because participants reported flash number, only the visual estimate entered the response stage. The three decision rules were model averaging, which weighted the common- and separate-cause estimates by their posterior probabilities; model selection, which selected the common-cause estimate when *P*(*C* = 1 | *x*, *t*) > 0.5; and probability matching, which sampled the causal structure according to this posterior. For model averaging, the reported visual estimate was

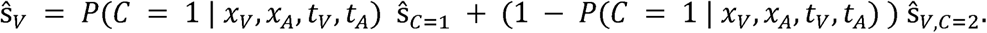

The continuous visual estimate was converted to the task response by reporting two flashes when it was at least 1.5 and one flash otherwise. The model prediction for each stimulus condition was therefore the probability of a two-flash response. In the primary, Standard 2D BCI model, the free parameters were *p_common_*, σ*_V_,* σ*_t_*, and δ. Auditory numerosity uncertainty and the numerosity-prior parameters were fixed (σ*_A_* = 0.33, σ*_p_* = 1.2, μ*_p_* = 1.4) because the task contained no unisensory auditory report with which to identify σ*_A_* independently and because freeing weakly constrained prior parameters can trade off against *p_common_*. Full evidence integrals, estimator equations, and identifiability and recovery analyses will be reported in the Supplementary Methods and Figure S1.

#### 2.5.2. Bayesian causal-inference model of simultaneity judgments

The SJ data were fitted with a temporal BCI model that used the same causal hypotheses and the same temporal parameterization as the SIFI model. On each trial, visual and auditory onset measurements were generated according to the temporal likelihoods above, with one shared σ*_t_* and an auditory–visual latency offset δ. Under *C* = 1, the two measurements were generated by one latent event time drawn from 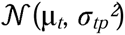; under *C* = 2, they were generated by independent latent event times drawn from the same prior. The posterior probability of simultaneity-relevant common causation was

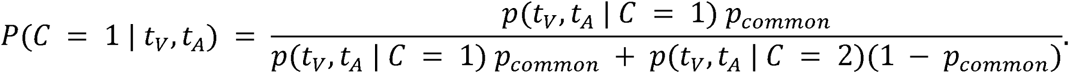

Thus, *p_common_* controlled the prior tendency to judge an audiovisual pair as arising from one event, σ*_t_* controlled the precision of temporal evidence, and δ translated the entire predicted simultaneity function along the SOA axis. These three parameters were estimated separately for every participant at pre- and post-test, whereas μ*_t_* = 0 ms and σ*_tp_* = 300 ms were fixed. We compared model selection, in which a “simultaneous” response was generated when *P*(*C* = 1 | *t_V_*, *t_A_*) > 0.5, with probability matching, in which the response was sampled from this posterior. A fixed lapse mixture (lapse rate = 0.02; simultaneous-response probability on lapse trials = 0.5) prevented occasional stimulus-independent errors from distorting the causal and temporal parameters. The temporal likelihoods and predicted response probabilities were evaluated deterministically by numerical quadrature; the equivalent log-space implementation and its analytic reduction will be provided in the Supplementary Methods.

#### 2.5.3. Model fitting and comparison

SIFI and SJ models were fitted separately to each participant and time point using the binomial likelihood of the observed response counts. SIFI probabilities were approximated from 10,000 simulated trials per condition using common random numbers so that the objective function remained deterministic across optimizer evaluations. Parameters were optimized with Bayesian Adaptive Direct Search using 20 randomized restarts per decision rule. The SJ likelihood was evaluated without Monte Carlo sampling and was fitted using the same optimizer and restart procedure. Within each architecture, the decision rule with the lowest Bayesian information criterion (BIC) was retained; BIC was calculated as *k* ln(*N*) + 2NLL, where *k* is the number of free parameters, *N* is the number of valid trials, and NLL is the minimized negative log-likelihood. Two-fold cross-validation, formed by alternating trials within each SIFI condition, provided an additional out-of-sample check.

The Standard 2D BCI model was compared with four theoretically targeted alternatives. The cross-task latency-constrained model fixed δ to the participant’s independently fitted SJ value while estimating σ*_t_* from SIFI and allowing auditory as well as visual numerosity uncertainty to vary. The fully flexible sensory–temporal model estimated both σ*_t_* and δ from SIFI and additionally allowed σ*_A_* to vary. The fixed-temporal constrained-numerosity model imposed σ*_t_* = 60 ms and δ = 0 while fixing σ*_A_,* whereas the fixed-temporal flexible-numerosity model retained this temporal channel but estimated σ*_A_*. Together, these alternatives tested whether fit improvements required individual temporal precision, a latency shift, free auditory precision, or a flexible numerosity prior. BIC comparisons used matched participant–time cells; full specifications and recovery analyses will be reported in the Supplementary Methods.

The theoretically focal computational outcomes were the SIFI common-cause prior (*p_common_*) and temporal uncertainty (σ*_t_*). Group × Time effects were evaluated by comparing participant-level post-minus-pre change scores between groups using two-sided Welch tests, with Holm correction across these two focal interactions. Following significant interactions, the four planned within-group simple effects arising from the two parameters and two groups were evaluated using two-sided Wilcoxon signed-rank tests and Holm-adjusted as a separate family. The SIFI sensory-noise ratio and all SJ parameter analyses were treated as secondary analyses, with their correction families specified in the Supplementary Methods.

### 2.6. Software and code

Bayesian causal inference models were implemented and fitted in MATLAB (version ⟨R2025b⟩; MathWorks, Natick, MA). Model code refers to Zhu et al., 2024b, BCI Toolbox. Parameter estimation used Bayesian Adaptive Direct Search (BADS, Acerbi and Ma, 2017; https://github.com/acerbilab/bads. Behavioural analyses, statistical tests and figures were produced in Python (version ⟨3.7⟩; with NumPy, SciPy, pandas and Matplotlib. All analysis and modelling code, together with the trial-level data, is available at https://github.com/evans1112/ADHD_perceptual_learning.

## 3. Results

### 3.1. Experiment 1

Children with ADHD showed greater susceptibility to the sound-induced fission illusion than typically developing (TD) children (Figure 1B). The mean fission-illusion rate was 0.857 (SD = 0.091) in the ADHD group and 0.771 (SD = 0.182) in the TD group, corresponding to a mean group difference of 0.078 (95% CI [0.009, 0.164]). This difference was significant in both a Welch’s t-test, *t*(39.63) = 2.25, *p* = 0.030, Cohen’s *d* = 0.60, and a two-sided Mann–Whitney test, *U* = 532, *p* = 0.021.

**Figure 1.**
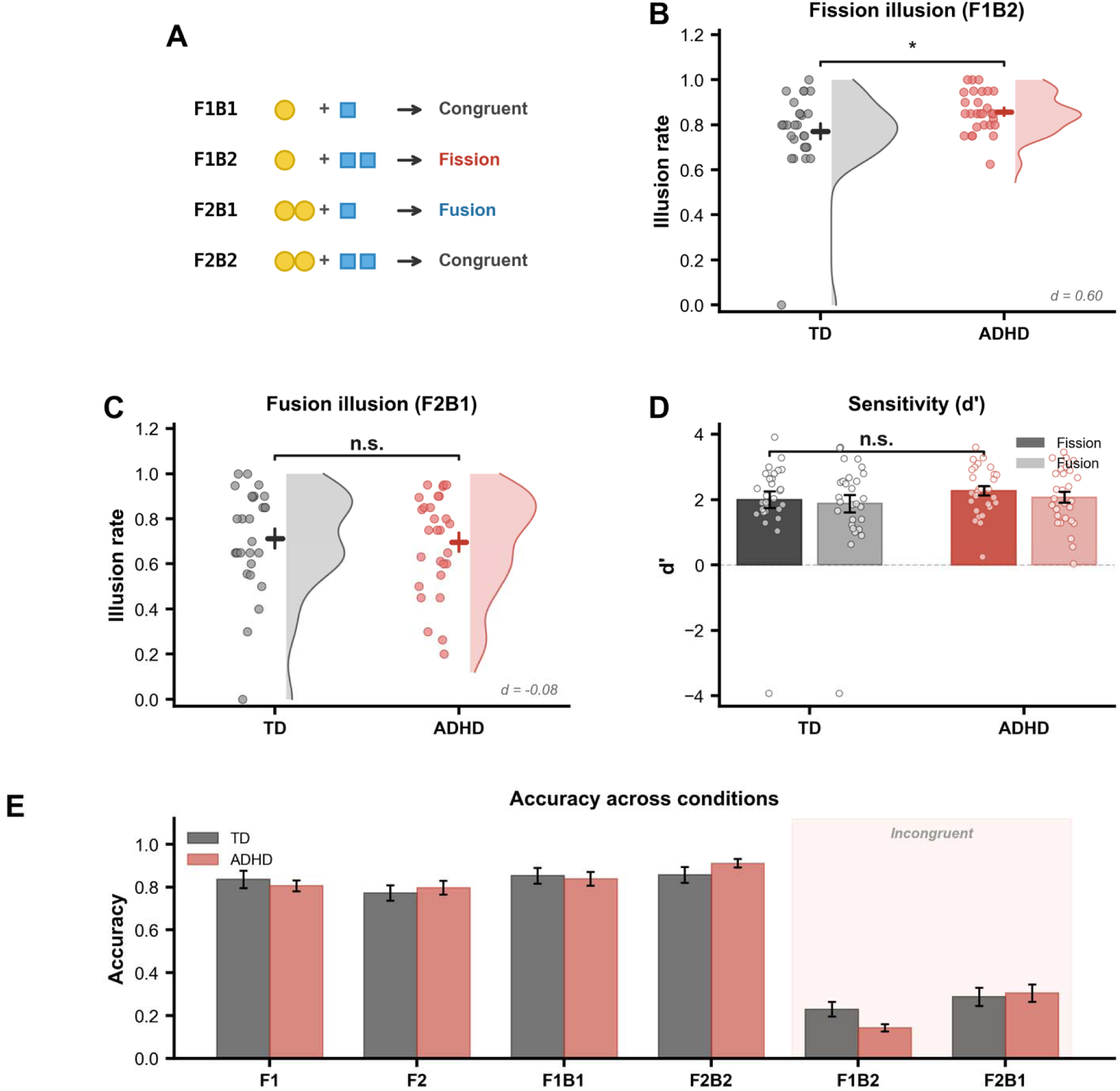
Children with ADHD showed enhanced susceptibility to the sound-induced fission illusion. **(A)** Schematic of the audiovisual stimulus conditions. Yellow circles represent visual flashes and blue squares represent auditory beeps. Congruent conditions comprised one flash paired with one beep (F1B1) and two flashes paired with two beeps (F2B2). Incongruent conditions comprised one flash paired with two beeps, eliciting the fission illusion (F1B2), and two flashes paired with one beep, eliciting the fusion illusion (F2B1). **(B)** Children with ADHD showed a higher fission-illusion rate than typically developing (TD) children. **(C)** Fusion-illusion rates did not differ between groups. In **(B–C)**, dots represent individual participants, horizontal lines and vertical bars denote the mean ± SEM, and half-violin plots show the distributions. Illusion rates were calculated across valid-response trials. **(D)** Signal-detection sensitivity (*d*′), calculated by contrasting each incongruent illusion condition with its corresponding congruent control condition, did not differ between groups for either or fusion. Bars and error bars indicate the mean ± SEM, with dots showing individual participants. **(E)** Mean response accuracy across unisensory, audiovisually congruent, and audiovisually incongruent conditions. No group differences were detected in the unisensory or congruent control conditions (all uncorrected *p* > .05). Error bars indicate SEM. TD, typically developing; ADHD, attention-deficit/hyperactivity disorder. *n* = 28 participants per group. Asterisks indicate *p* < 0.05; n.s., not significant.

This group difference was not observed for the fusion illusion (Figure 1C). Fusion-illusion rates were comparable between children with ADHD (M = 0.695, SD = 0.218) and TD children (M = 0.712, SD = 0.228), *t*(53.88) = −0.29, *p* = 0.774, mean difference = −0.017, 95% CI [−0.137, 0.102], *d* = −0.08. A direct comparison of the group effects across illusion types yielded a modest Group × Illusion-type interaction, *t*(51.80) = 1.95, *p* = 0.056, with the fission–fusion contrast being 0.104 larger in the ADHD group (95% CI [-0.003, 0.211]). Thus, the group difference was concentrated in the fission condition rather than reflecting a general increase in susceptibility to audiovisual illusions.

Signal-detection analyses did not indicate corresponding group differences in illusion-specific sensitivity (Figure 1D). Fission sensitivity was similar in children with ADHD (M = 2.27, SD = 0.74) and TD children (M = 2.00, SD = 1.34), *t*(42.26) = 0.94, *p* = 0.352. Fusion sensitivity was also comparable between groups (ADHD: M = 2.08, SD = 0.89; TD: M = 1.88, SD = 1.41), *t*(45.70) = 0.61, *p* = 0.544. Accuracy in the unisensory and audiovisually congruent control conditions did not differ between groups (all uncorrected *p* ≥ 0.202; all Holm-adjusted *p* = 1.000; Figure 1E), arguing against a generalized impairment in visual-number discrimination or task performance.

An exploratory response-criterion analysis suggested a lower fission criterion in the ADHD group (ADHD: M = −0.016; TD: M = 0.187), Welch’s *t*(51.88) = −2.08, *p* = 0.043. However, this difference was not significant in the corresponding nonparametric analysis, *U* = 282.5, *p* = 0.074, and should therefore be interpreted cautiously.

### 3.2. Experiment 2

Fission-illusion reports varied markedly as a function of audiovisual stimulus onset asynchrony (SOA; Figure 2B). A mixed-design analysis with Group as a between-participant factor and SOA as a within-participant factor revealed a strong main effect of SOA, Greenhouse–Geisser-corrected *F*(4.21, 243.94) = 27.34, *p* = 1.97 × 10^−19^, but no main effect of Group, *F*(1, 58) = 0.77, *p* = 0.384. Averaged across SOAs, illusion rates were therefore comparable between children with ADHD (M = 0.501, SD = 0.255) and typically developing (TD) children (M = 0.450, SD = 0.186), *t*(53.10) = 0.88, *p* = 0.384, mean difference = 0.051, 95% CI [−0.065, 0.166], Cohen’s *d* = 0.23.

**Figure 2.**
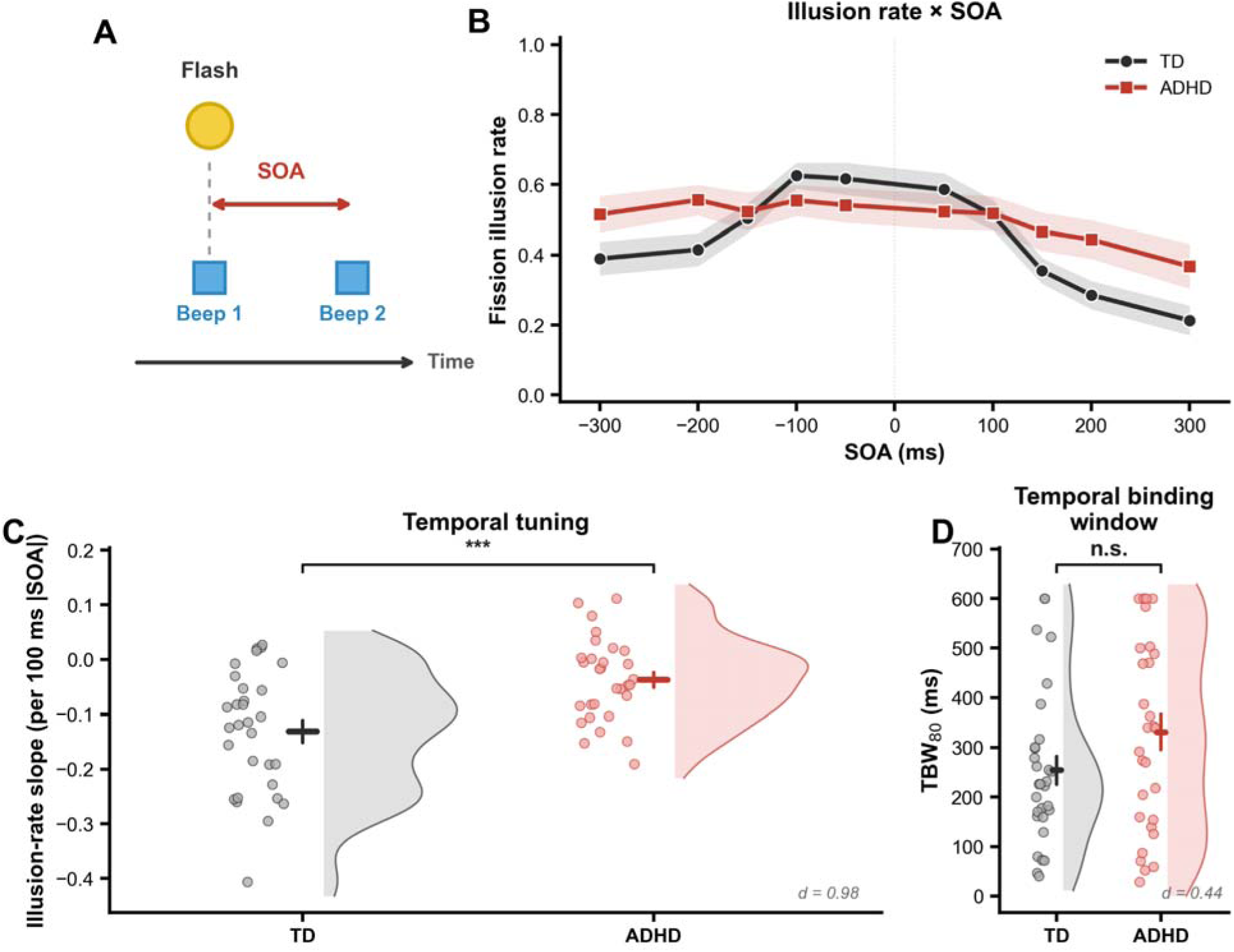
Children with ADHD showed reduced temporal selectivity of audiovisual binding without a significant increase in conventional temporal-binding-window width. **(A)** Schematic of the F1B2 sound-induced flash illusion condition used to characterize temporal binding. Yellow circles represent visual flashes and blue squares represent auditory beeps; the audiovisual stimulus onset asynchrony (SOA) was manipulated across trials. **(B)** Mean fission-illusion rate as a function of SOA in typically developing (TD) children and children with ADHD. Negative SOAs indicate that the auditory stimulus led the visual stimulus, whereas positive SOAs indicate that the visual stimulus led. Shaded regions denote SEM. A mixed-design analysis revealed a strong main effect of SOA, Greenhouse–Geisser-corrected *F*(4.21, 243.94) = 27.34, *p* = 1.97 × 10^−19, and a significant Group × SOA interaction, *F*(4.21, 243.94) = 6.96, *p* = 1.76 × 10^−5, but no main effect of Group, *F*(1, 58) = 0.77, *p* = 0.384. No individual SOA-level group comparison survived Holm correction. **(C)** Individual temporal-tuning slopes, calculated as the linear change in illusion rate for each 100-ms increase in absolute SOA. More negative values indicate stronger attenuation of the illusion with increasing audiovisual asynchrony. Slopes were shallower in children with ADHD than in TD children, indicating reduced temporal selectivity, Welch’s *t*(50.84) = 3.81, *p* = 0.0004, mean difference = 0.093, 95% CI [0.044, 0.142], Cohen’s *d* = 0.98. **(D)** Individual nonparametric temporal-binding-window estimates (TBW_₈₀_), defined as the interpolated SOA range over which the illusion rate remained at or above 80% of the participant-specific maximum. TBW_₈₀_ was numerically larger in children with ADHD but did not differ significantly between groups, *t*(55.19) = 1.70, *p*= 0.096, mean difference = 77.03 ms, 95% CI [−13.99, 168.06], *d* = 0.44. In **(C–D)**, dots represent individual participants, horizontal and vertical lines indicate the mean ± SEM, and half-violin plots show the distributions. Illusion rates were calculated across valid-response trials. *n* = 30 participants per group. *** indicates *p* < 0.001; n.s., not significant.

Critically, the effect of SOA differed between groups, as indicated by a significant Group × SOA interaction, Greenhouse–Geisser-corrected *F*(4.21, 243.94) = 6.96, *p* = 1.76 × 10^−5^. In TD children, illusion reports decreased substantially as the absolute audiovisual asynchrony increased, whereas the ADHD group showed a considerably flatter temporal response profile. At near SOAs (|SOA| ≤ 100 ms), mean illusion rates were 0.585 in TD children and 0.534 in children with ADHD. At far SOAs (|SOA| ≥ 200 ms), illusion rates decreased to 0.326 in TD children but remained at 0.470 in the ADHD group. Consequently, the far-minus-near attenuation was substantially smaller in ADHD (M = −0.064, SD = 0.142) than in TD children (M = −0.260, SD = 0.221), *t*(49.52) = 4.07, *p* = 0.0002, mean difference = 0.196, 95% CI [0.099, 0.292], *d* = 1.05.

The same pattern was observed when temporal selectivity was quantified for each participant as the slope of illusion rate over absolute SOA. Slopes were strongly negative in TD children (M = −0.131 per 100 ms, SD = 0.111), indicating progressive attenuation of the illusion with increasing asynchrony, but were considerably shallower in children with ADHD (M = −0.038, SD = 0.075; Figure 2C). The group difference was significant, *t*(50.84) = 3.81, *p* = 0.0004, mean difference = 0.093, 95% CI [0.044, 0.142], *d* = 0.98, and was corroborated by a two-sided Mann–Whitney test, *U* = 668, *p* = 0.0013. None of the individual SOA-level group comparisons survived Holm correction (all adjusted *p* ≥ 0.242), indicating that the interaction reflected a distributed difference in the shape of the temporal response function rather than an isolated effect at a single SOA.

In contrast, conventional estimates of temporal-binding-window width did not differ significantly between groups. The nonparametric TBW₈₀ estimate was numerically larger in children with ADHD (M = 330.83 ms, SD = 194.62) than in TD children (M = 253.80 ms, SD = 154.99; Figure 2D), but the 77.03-ms difference was not significant in a two-sided test, *t*(55.19) = 1.70, *p* = 0.096, 95% CI [−13.99, 168.06], *d* = 0.44. Gaussian width estimates were also indistinguishable between groups (ADHD: M = 170.88 ms, SD = 134.48; TD: M = 177.70 ms, SD = 123.37), *t*(57.60) = −0.20, *p* = 0.839, *d* = −0.05.

Notably, Gaussian fits provided a poorer description of the ADHD response profiles (M *R*² = 0.581, SD = 0.232) than of the TD profiles (M *R*² = 0.735, SD = 0.226), *t*(57.96) = −2.59, *p* = 0.012. This difference is consistent with the relatively elevated illusion rates at long SOAs in ADHD and indicates that the group difference is not well characterized as a uniform increase in Gaussian width. When illusion rates were compared with the F1B1 control condition, the illusion remained significant at +300 ms in the ADHD group (Holm-adjusted *p* = 0.0006), whereas the corresponding effect in the TD group did not reach significance (adjusted *p* = 0.058). This within-group pattern is consistent with more persistent binding in ADHD but does not, by itself, constitute a direct test of the group difference.

No group difference in the control conditions survived Holm correction (all adjusted *p* ≥ 0.060), and missed-response rates were comparable between children with ADHD (M = 7.22%) and TD children (M = 6.10%), *t*(45.77) = 0.66, *p* = 0.512. Together, these results indicate that ADHD was associated with reduced temporal selectivity of audiovisual binding rather than a general increase in illusion susceptibility or a uniformly enlarged temporal binding window.

### 3.3. Experiment 3

#### 3.3.1. Behavioural results

Training reduced susceptibility to the sound-induced flash illusion in both groups (Figure 3A,D). In typically developing children, the mean fission-illusion rate decreased from 0.49 (SD = 0.27) before training to 0.33 (SD = 0.27) after training, mean change = −0.16, 95% CI [−0.29, −0.03], *t*(26) = −2.55, *p* = .017, *d*_z_ = −0.49. Children with ADHD showed a comparable decrease, from 0.54 (SD = 0.26) to 0.40 (SD = 0.20), mean change = −0.14, 95% CI [−0.25, −0.03], *t*(22) = −2.61, *p* = .016, *d*_z_ = −0.54. Both effects remained significant after Holm correction within this endpoint (both adjusted *p* < .05). The Group × Time interaction was not significant, *t*(47.79) = 0.23, *p* = .82, indicating that the overt reduction in illusion susceptibility was similar across groups.

**Figure 3.**
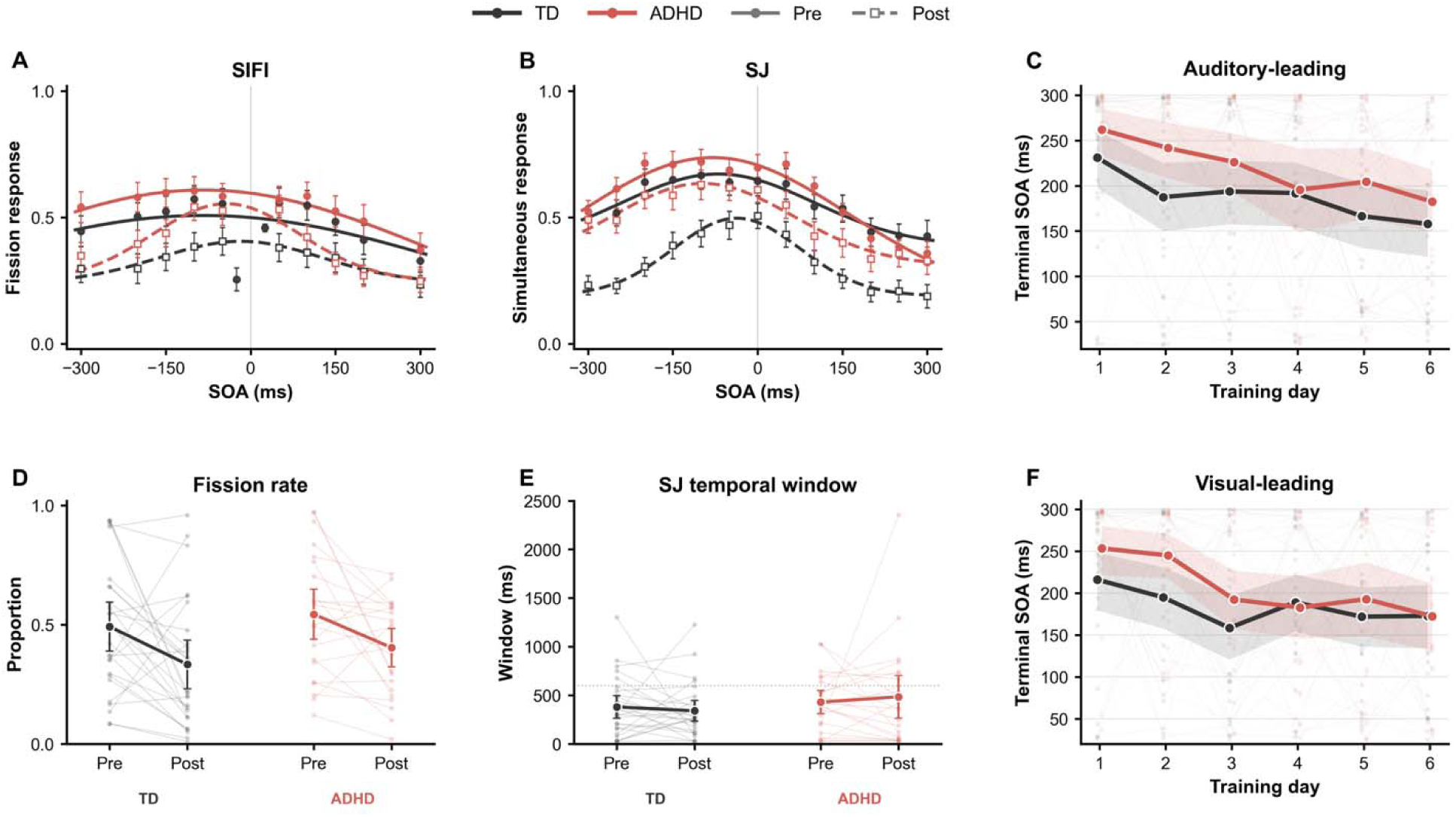
Training progression and pre–post changes in audiovisual temporal judgments. (A) Sound-induced flash illusion (SIFI) curves showing the proportion of fission responses as a function of stimulus-onset asynchrony (SOA) before and after training. (B) Simultaneity-judgment (SJ) curves showing the proportion of trials judged simultaneous. In A and B, symbols show group means ± SEM at each SOA, and smooth lines show functions fitted to the participant-averaged psychometric data. Negative SOAs indicate auditory-leading trials and positive SOAs indicate visual-leading trials. (C) Daily progression of the auditory-leading adaptive staircase across the six training sessions. (D) Mean fission-illusion rate across SIFI SOAs for each participant before and after training. (E) Individual SJ temporal windows, defined as the full width at half maximum of each participant’s fitted simultaneity curve. The horizontal dotted line indicates the 600-ms tested SOA span; estimates above this value depend on extrapolation beyond the observed stimulus range. (F) Daily progression of the visual-leading adaptive staircase. In C and F, terminal staircase level was defined as the mean absolute SOA of the final 20 non-baseline adaptive trials within each session and temporal direction. Small points and faint lines represent individual participants, thick lines and large points represent group means, and shaded bands indicate participant-bootstrap 95% confidence intervals. Lower terminal SOAs indicate progression to more difficult temporal offsets but should not be interpreted as independent psychophysical threshold estimates. In D and E, small points and thin lines represent individual participants, large points represent group means, and error bars indicate 95% confidence intervals. TD participants are shown in black and participants with ADHD in red.

The trained simultaneity-judgment task showed a more differentiated pattern (Figure 3B,E). In the TD group, the mean proportion of simultaneous responses decreased substantially from 0.56 before training to 0.32 after training, mean change = −0.24, 95% CI [−0.36, −0.12], *t*(26) = −4.06, *p* < .001, *d*_z_ = −0.78. This effect remained significant after Holm correction, adjusted *p* = .001. In the ADHD group, simultaneous responses decreased from 0.59 to 0.50, but this change did not reach significance, mean change = −0.09, 95% CI [−0.20, 0.01], *t*(22) = −1.94, *p* = .065, *d*_z_ = −0.41. The nominal Group × Time interaction, *t*(47.55) = 1.87, *p* = .067, did not survive correction for multiple comparisons, adjusted *p* = .13. Thus, training produced a particularly marked reduction in the tendency to classify audiovisual events as simultaneous in TD children, although the evidence for a group difference was not conclusive after correction.

At the individual level, fitted SJ temporal windows did not change reliably in either group (Figure 3E). In TD children, the mean temporal window decreased from 382 ms (SD = 306) to 341 ms (SD = 279), mean change = −40 ms, 95% CI [−193, 113], *t*(26) = −0.54, *p* = .593, *d*_z_ = −0.10. In children with ADHD, the mean window changed from 430 ms (SD = 291) to 485 ms (SD = 533), mean change = 55 ms, 95% CI [−191, 301], *t*(22) = 0.46, *p* = .648, *d*_z_ = 0.1. The Group × Time interaction was also not significant, *t*(37.81) = 0.68, *p* = .501. Training therefore altered the overall probability of simultaneous responding, particularly in TD children, but conventional individual-level SJ window estimates provided no reliable evidence of temporal-window narrowing.

Complementary paired-bootstrap fits to the participant-averaged curves showed numerical narrowing of the SJ function in both groups (TD: Δ*window* = −136 ms, *p_boot_*=.117; ADHD: Δ*window* = −190 ms, *p_boot_*=.019), but the magnitude of narrowing did not differ between groups (ADHD-minus-TD difference = −54 ms, 95% CI [−315, 221], *p_boot_* =.659). This group-curve effect should be distinguished from the individual-window endpoint, which provided no evidence that temporal narrowing occurred consistently across participants.

Temporal-order judgments showed no corresponding improvement in temporal resolution (Figure S2). Accuracy increased numerically from 0.57 to 0.58 in TD children, but this change was not significant, mean change = 0.02, 95% CI [−0.04, 0.07], *t*(26) = 0.63, *p* = .532, *d*_z_ = 0.12. Accuracy was essentially unchanged in children with ADHD, decreasing from 0.58 to 0.58, mean change = −0.004, 95% CI [−0.04, 0.03], *t*(22) = −0.20, *p* = .842, *d*_z_ = −0.04. The Group × Time interaction was not significant, *t*(45.48) = −0.63, *p* = .531.

Together, the behavioral results demonstrate that training reduced SIFI susceptibility in both groups and strongly altered simultaneity judgments in TD children, but did not produce reliable narrowing of conventional individual-level SJ or TOJ temporal windows. The absence of a clear behavioral-window effect is important for interpreting the computational analyses: any training-related change in latent temporal precision identified by the model cannot be treated as a simple restatement of the conventional TBW results, but instead reflects information recovered by decomposing behavior into distinct inferential components.

#### 3.3.2. Computational modelling

Before considering training effects, we compared baseline (pre-training) parameter estimates between groups. In the SIFI model, temporal uncertainty was higher in children with ADHD (M = 228.86 ms, SD = 170.58) than in TD children (M = 114.10 ms, SD = 140.11), *t*(42.64) = 2.57, *p* = .014, mean difference = 114.77 ms, 95% CI [24.73,204.80]). No baseline group differences were detected for *p_common_*, ***σ****_V_*, or (all *p* > 0.05; See *Methods* for definitions).

We next asked whether the behavioral effects of training reflected changes in temporal evidence, causal expectations, or lower-level sensory uncertainty. The two theoretically focal SIFI parameters were the common-cause prior (*p_common_*)and temporal uncertainty (***σ****_t_*). Their Group × Time effects were evaluated as a two-test family using Holm correction. Because both interactions survived correction, the four corresponding within-group simple effects were examined as a separate Holm-corrected family. Analyses of the SIFI sensory-noise ratio and all SJ parameters were treated as secondary.

For the simultaneity-judgment task, the estimated common-cause prior decreased numerically in both groups (Figure 4A). In the TD group, *p_common_* decreased from 0.586 (SD = 0.245) before training to 0.399 (SD = 0.177) after training (mean change = −0.186, 95% CI [−0.316, −0.057], *p* =.009, *p_Holm_* =.053). The corresponding decrease in the ADHD group was smaller (0.633 to 0.549; mean change = −0.083, 95% CI [−0.183, 0.017], *p_Holm_* =.117). However, the difference in *p_common_* change between groups was not significant after correction, (*t*(46.45)=1.30, *p_Holm_* =.398). Neither group showed reliable changes in temporal noise, (***σ****_t_*) (Figure 4B; TD: *p_Holm_* =.679; ADHD: *p_Holm_* =.808), or in the latency-offset parameter, (δ) (Figure 4C; both adjusted *p_Holm_* = .808). The group differences in change were also nonsignificant for both parameters (***σ****_t_*: *p_Holm_* =.310; : *p_Holm_*=.398). In this secondary analysis of the trained task, neither within-group reduction in *p*_common_ survived correction, although the direction of change was consistent with the focal SIFI result.

**Figure 4.**
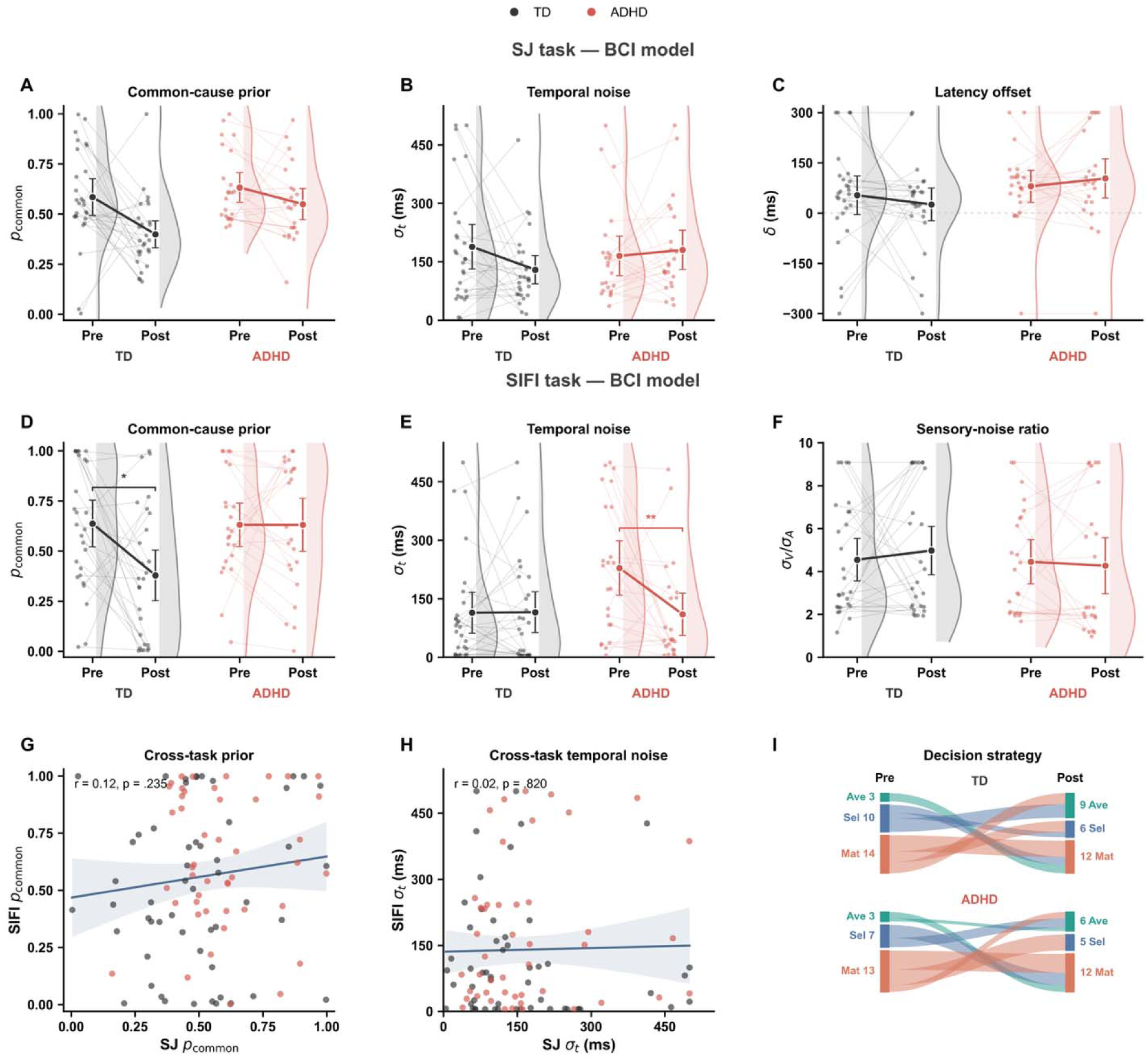
Computational modelling reveals dissociable training-related changes in causal priors and temporal precision. (A–C) Parameter estimates from the Bayesian causal-inference (BCI) model fitted to the simultaneity-judgment (SJ) task, showing the common-cause prior (*p_common_*; A), temporal noise ( *_t_*; B), and latency offset ( ; C) before and after training. (D–F) Corresponding parameter estimates from the BCI model fitted to the sound-induced flash illusion (SIFI), showing *p_common_* (D), *_t_* (E), and the visual-to-auditory sensory-noise ratio (F). TD children showed a reduction in *p_common_*in the SIFI task, with a numerically consistent but non-significant reduction in the SJ task, whereas children with ADHD showed a selective reduction in SIFI *_t_*, indicating increased temporal precision. Small points represent individual participants, thin lines connect repeated observations, and half-violin distributions show the estimated parameter distributions. Large points and error bars indicate group means and 95% confidence intervals. Asterisks denote significant within-group pre–post changes assessed using Wilcoxon signed-rank tests with Holm correction within each task (\**p* <.05), (\*\**p* <.01). (G–H) Cross-task associations between SJ and SIFI estimates of *p_common_* (G) and *_t_* (H), pooling pre- and post-training estimates as 100 observations. Lines show Pearson regression fits and shaded regions indicate 95% confidence intervals. Neither parameter showed a significant cross-task correlation. (I) Alluvial plots showing transitions among model averaging (Ave), model selection (Sel), and probability matching (Mat) strategies from pre- to post-training. Flow widths and adjacent numbers represent participant counts. Strategy switching was frequent at the individual level but did not produce a significant group-level redistribution of strategies in either group.

A clearer pattern emerged when the BCI model was applied to the sound-induced flash illusion (Figure 4D–F). The Standard 2D BCI model showed an overall mean R^2^ of .751 across evaluable participants. Consistent with the SJ estimates, TD children showed a substantial reduction in *p_common_*, from 0.637 (SD = 0.309) to 0.379 (SD = 0.335; mean change = −0.259, 95% CI [−0.435, −0.082], *p_Holm_* =.021); Figure 4D). In contrast, *p_common_* remained stable in the ADHD group (0.631 to 0.630; mean change = −0.001, 95% CI [−0.145, 0.143], *p_Holm_* =1.000). The group difference in change was significant, *t*(47.23)=2.33, *p*=.024, *p_Holm_*=.048.

Temporal noise showed the complementary pattern (Figure 4E). In children with ADHD, ***σ****_t_* decreased from 228.86 ms (SD = 170.58) to 110.19 ms (SD = 132.04), corresponding to a mean reduction of 118.67 ms (95% CI [32.78, 204.56], *p_Holm_* =.006). Because lower ***σ****_t_* indicates more precise temporal evidence, this change reflects a substantial improvement in audiovisual temporal precision. No corresponding change was observed in TD children (114.10 to 115.49 ms; mean change = 1.39 ms, 95% CI [−63.37, 66.26], *p* =1.000). Critically, the group difference in temporal-noise change was significant, (t(42.76)=-2.31, *p* =.026, *p_Holm_*=.048). By contrast, the visual-to-auditory sensory-noise ratio, (**σ***_V_*/***σ****_A_*), remained stable in both groups (Figure 4F; TD: *p* =1.000; ADHD: p =1.000), with no group difference in change, (*t*(46.93)=-0.63), *p* =.532). The ADHD-related reduction in ***σ****_t_* therefore cannot be explained by a general change in relative unisensory noise.

We then examined whether individual parameter estimates generalized across the SJ and SIFI tasks. Following the planned analysis, pre- and post-training observations were pooled and treated as 100 observations. There was no reliable cross-task association for either the common-cause prior, (*r*=.12, 95% CI [−.08, .31], *p* =.235) (Figure 4G), or temporal noise, (*r*=.02, 95% CI [−.17, .22], *p* =.820) (Figure 4H). Thus, although the group-level training effects converged on interpretable computational components, individual estimates of these components did not show evidence of task-general correspondence.

Finally, we examined whether training altered the decision strategy inferred from the SIFI model—model averaging, model selection, or probability matching (Figure 4I). Individual strategy switching was common: 19 of 27 TD children (70.4%) and 16 of 23 children with ADHD (69.6%) changed their assigned strategy between sessions. Nevertheless, the marginal distribution of strategies did not change systematically in either group (TD: Stuart–Maxwell □^2^(2)=3.23, *p*=.199; ADHD: □^2^(2) =1.33, *p* =.513). The continuous parameter effects therefore occurred without a reliable group-level redistribution among discrete decision strategies.

Together, these analyses reveal two dissociable computational expressions of learning. Both focal Group × Time effects survived Holm correction across the two primary SIFI parameters: TD children showed a selective reduction in the prior probability assigned to a common audiovisual cause, whereas children with ADHD showed a selective increase in the precision of temporal evidence. Training-related behavioral improvement therefore did not arise from a single common mechanism across groups.

#### 3.3.3. Model Comparison

Candidate architectures differed reliably in their BIC ranks across the 100 matched participant– time cells (Friedman ^2^(4)=72.82), *p* < 0.001, Kendall’s *W* =.182; Figure 5A). The Standard 2D BCI model was the most frequent winner, producing the lowest BIC in 50% of the cells. By comparison, the cross-task latency-constrained model was preferred in 12 cells (12%), the fully flexible sensory–temporal model in none, the fixed-temporal constrained-numerosity model in 16 cells (16%), and the fixed-temporal flexible-numerosity model in 22 cells (22%).

**Figure 5.**
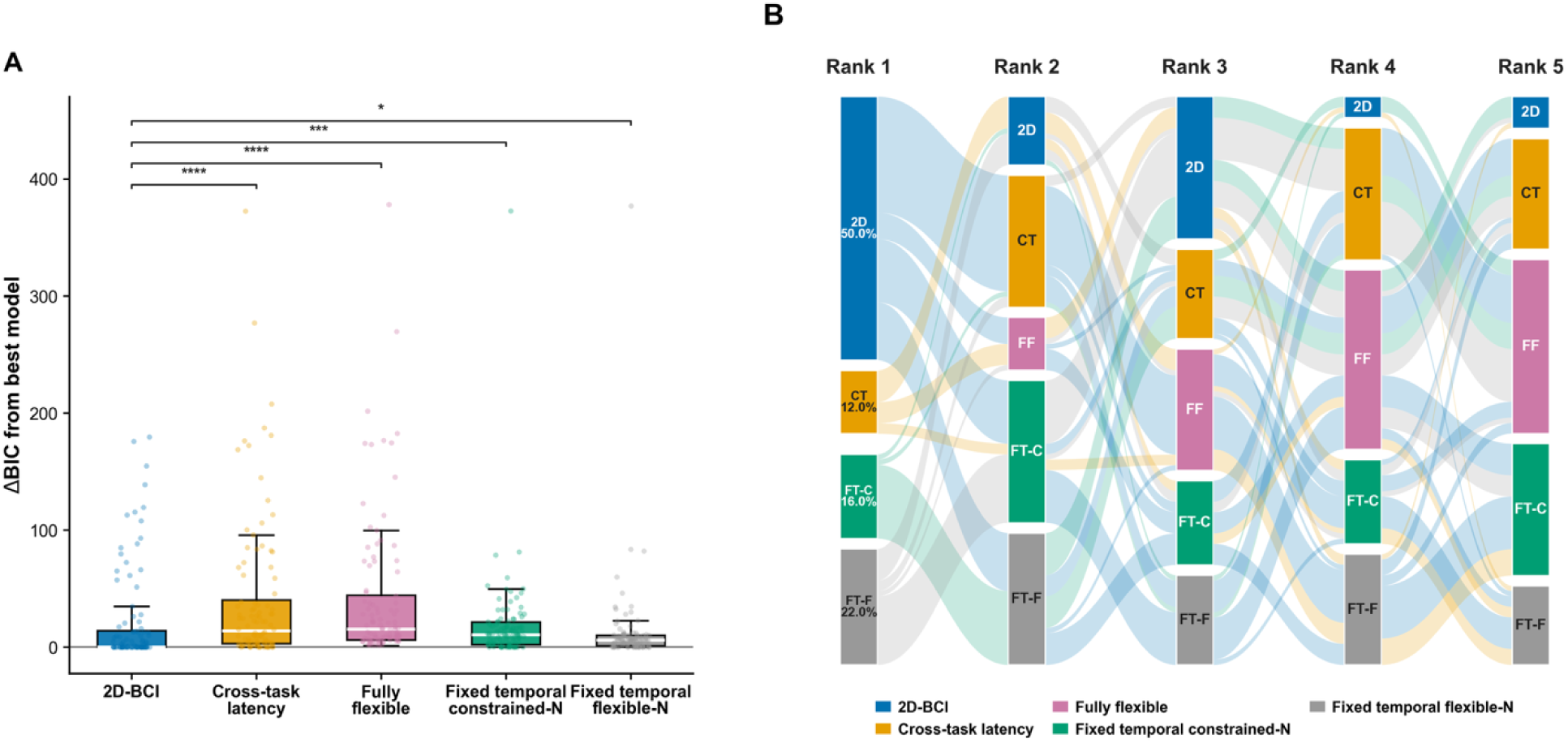
Comparison of candidate Bayesian causal inference architectures. **(A)** Distribution of ΔBIC across participants, calculated relative to the lowest-BIC model within each cell; lower values therefore indicate stronger relative support. Points represent individual cells, boxes show the interquartile range with the median indicated by the white line. Brackets show planned comparisons between the Standard 2D BCI model and each alternative, evaluated using exact paired sign tests based only on which model produced the lower BIC. p-values were Holm-corrected across the four comparisons (\**p_Holm_*<.05, \*\**p_Holm_*<.01, \*\*\**p_Holm_*<.001, \*\*\*\**p_Holm_*<.0001). **(B)** Alluvial representation of the complete model-ranking distribution. Models were ranked from the lowest BIC (Rank 1) to the highest BIC (Rank 5) within each participant–time cell. Node height and flow width represent the proportion of cells following each ranking path; flow colors are inherited from the Rank 1 model, allowing each winning-model cohort to be traced through subsequent ranks. Percentages within Rank 1 nodes indicate the proportion of cells in which each model achieved the lowest BIC. 2D-BCI, Standard 2D Bayesian causal-inference model; CT, cross-task latency-constrained model; FF, fully flexible sensory–temporal model; FT-C, fixed-temporal constrained-numerosity model; FT-F, fixed-temporal flexible-numerosity model.

Planned pairwise comparisons evaluated only the direction of the within-cell BIC difference, that is, which model yielded the lower BIC, using exact sign tests with Holm correction across the four comparisons. The Standard 2D BCI model was favored over the cross-task latency-constrained model in 78 versus 22 cells (*p_Holm_*=4.77*10^-8^), over the fully flexible sensory– temporal model in 88 versus 12 cells (*p_Holm_*=7.65*10^-15^), over the fixed-temporal constrained-numerosity model in 62 versus 38 cells (*p_Holm_*=.021), and over the fixed-temporal flexible-numerosity model in 67 versus 45 cells (*p_Holm_*=.0467). The full ranking distribution reinforced this pattern: the Standard 2D BCI model occupied one of the top three positions in 90% of the cells (Figure 5B). Thus, the focal model was the most consistently supported architecture across participant, although model preference remained heterogeneous, with an alternative model ranking first in 50% of cells.

## 4. Discussion

Perceptual learning brought children with and without ADHD to a similar behavioral endpoint, but it did not take them there by the same route. After simultaneity-judgment training, both groups became less susceptible to the sound-induced flash illusion (SIFI). Beneath this convergence, however, the Bayesian causal inference model revealed an asymmetric pattern: learning in typically developing (TD) children was expressed primarily as a reduction in the causal prior that auditory and visual signals share a common cause (*p_common_*), whereas learning in children with ADHD was expressed as a reduction in temporal uncertainty (***σ****_t_*) within the SIFI model. Unisensory numerosity precision and the distribution of decision strategies showed no corresponding systematic shift. The same perceptual change can therefore conceal different forms of plasticity, one that alters what the observer expects about the causal structure of the world, and another that alters how precisely the observer represents the temporal evidence arriving from it.

This distinction changes how multisensory perceptual learning should be interpreted. A reduction in audiovisual binding is often treated as evidence that a single temporal binding window has narrowed (Powers et al., 2009; Wallace and Stevenson, 2014). Within Bayesian causal inference, however, binding depends jointly on the multidimensional likelihoods, the prior probability of a common source, and the strategy used to translate the resulting posterior into a response (Körding et al., 2007; Wozny et al., 2010; Rohe and Noppeney, 2015). Lowering *p_common_* makes integration less likely before the current temporal evidence is considered; lowering ***σ****_t_* makes an observed asynchrony more diagnostic of separate causes. Either operation can reduce illusion susceptibility. The present pattern is thus an instance of computational equifinality: behavior is many-to-one with respect to its latent causes. This is precisely why the comparable pre–post reduction in SIFI susceptibility does not imply equivalent learning across groups.

The earlier experiments anticipated this conclusion without resolving it. ADHD was associated with selectively greater fission susceptibility in Experiment 1, whereas Experiment 2 showed an altered SOA-dependent illusion profile rather than a simple, uniform enlargement of a Gaussian temporal window. That distinction matters. SIFI susceptibility reflects not only temporal tolerance, but also numerosity sensory reliability, causal expectation, attention, stimulus context, and response strategy (Shams et al., 2002; Noppeney, 2021; Colonius and Diederich, 2025; Zhu et al., 2026a; Wang et al., 2025). More broadly, meta-analytic evidence indicates that timing in ADHD is both less accurate and more variable, particularly at subsecond scales, but also that the size and even direction of differences depend on the temporal operation being measured (Zheng et al., 2022; Marx et al., 2022). The three experiments together therefore argue against translating ‘more illusion’ directly into ‘a wider temporal binding window’. They instead locate the informative difference in how the system changes when it is asked to learn.

One explanation for the divergent trajectories is selective rigidity in causal-prior updating. The trained task repeatedly supplied evidence that small audiovisual discrepancies should be classified as asynchronous. TD children appear to have extracted a higher-order regularity from that experience and reduced their default tendency to bind. Children with ADHD updated the temporal likelihood but did not show the same reduction in *p_common_*. On this account, the relevant limitation is not an inability to learn, but a failure to propagate learning from improved sensory discrimination to a slower, more abstract estimate of environmental causal structure. Such an interpretation resonates with evidence of reduced dynamic brain reconfiguration in ADHD across default-mode, attention, executive, and primary networks (Yin et al., 2022), and with findings that children with ADHD occupy task-suboptimal brain states associated with variable behavior and slower evidence accumulation (Cai et al., 2021). It also offers a perceptual counterpart to cognitive accounts of impaired set shifting, while identifying a more specific target than “inflexibility” in general: the adjustment of a causal prior in response to accumulated multisensory evidence.

Yet prior rigidity should be treated as a candidate mechanism, not as a global description of the ADHD learner. Rapid visual statistical learning and the use of low-level perceptual priors can be intact in adults with ADHD (Richards et al., 2020). In reward learning, ADHD has also been associated with excessive, rather than insufficient, switching: Aster et al. (2024) found blunted reinforcement sensitivity, noisier choice changes, and enhanced learning from negative feedback. The apparent contradiction is theoretically useful. ADHD may not be characterized by uniformly low flexibility, but by a mismatch between levels and timescales of updating, which means too much influence from recent or salient events in some settings, yet insufficient revision of slowly accumulated contextual beliefs in others. In the present task, sharpening ***σ****_t_* can be driven by local trial-wise feedback, whereas changing *p_common_* requires evidence to be integrated across trials into a more stable model of how often signals belong together. A learner weighted toward immediate feedback could therefore improve the proximal temporal computation while leaving the slower causal statistic relatively unchanged. This proposal also connects naturally to classic accounts of ADHD emphasizing altered reinforcement timing and delay aversion (Sonuga-Barke, 2003; Yu and Sonuga-Barke, 2020).

A second, complementary account is that training repaired the computation with the greatest scope for improvement. Children with ADHD entered the SIFI assessment with greater temporal uncertainty (see Results), and Experiment 2 independently showed a flatter temporal modulation of the illusion. The large post-training reduction in their SIFI ***σ****_t_* may therefore reflect bottleneck-specific learning: feedback preferentially sharpened the dimension that was initially least stable. This interpretation fits the broader ADHD timing literature (Marx et al., 2022), but ***σ****_t_* should not be equated too literally with the width of a dedicated sensory neuron’s temporal tuning. In a behavioral observer model, it can also absorb trial-to-trial fluctuations in attention, arousal, encoding, and evidence quality. Elevated intra-individual variability is one of the most reproducible cognitive features of ADHD, and has been linked to unstable task engagement and transitions into suboptimal network states (Castellanos et al., 2009; Cai et al., 2021; Tamm et al., 2025). A lower post-training ***σ****_t_* could therefore mean that temporal representations became sharper, that attention to audiovisual onset became more stable, or both.

Another possibility concerns credit assignment. Perceptual learning requires the system to decide what caused its error: noisy timing, a biased sensory mapping, an inappropriate causal expectation, or a response rule. Multisensory causal inference itself can regulate recalibration by determining whether a discrepancy should be attributed to a common event and used as a teaching signal (Hong et al., 2021; Li et al., 2025). Attention further changes common-cause expectations and the gain assigned to prediction errors (Badde et al., 2020; Noppeney, 2021). TD and ADHD children may therefore have learned from the same feedback while assigning the error to different levels of the generative model. TD children behaved as if repeated errors changed the inferred statistics of the environment; ADHD children behaved as if those errors called for a more precise estimate of audiovisual timing. This need not reflect an explicit strategy. It could emerge from differences in attentional selection, confidence, error monitoring, or the relative precision assigned to top-down and bottom-up signals. Recent electrophysiological evidence provides a systems-level analogue of this possibility. Park et al. (2026) found that young people with the ADHD-inattentive presentation showed greater occupancy and longer dwell times in a posterior-to-anterior bottom-up mode, together with reduced expression of the opposing anterior-to-posterior top-down mode. Although these resting-state dynamics cannot be mapped directly onto likelihoods and priors, respectively, they provide a plausible neural context for the present asymmetry: training-related changes may be expressed more readily in the precision of incoming temporal evidence than in a higher-order causal expectation. Earlier ERP work, for example, found preserved early error detection but attenuated later error evaluation and post-error adjustment in children with ADHD (Wiersema et al., 2005).

The dissociation may also reflect the developmental organization of multisensory plasticity. Audiovisual temporal sensitivity and recalibration mature over a prolonged period extending into adolescence (Hillock-Dunn and Wallace, 2012; Rohlf et al., 2020; Ampollini et al., 2024; Wang et al., 2025). Integration can emerge before cumulative recalibration, indicating that using multisensory information online and updating the representations that support future inference are developmentally separable capacities (Rohlf et al., 2020). At the neural level, sensory estimates, causal-structure representations, and final perceptual decisions are distributed across a hierarchy involving sensory, superior temporal, parietal, and frontal regions (Powers et al., 2012; Rohe and Noppeney, 2015; Kayser and Shams, 2015). The present behavioral dissociation raises the possibility that training engages different parts of this hierarchy across groups: greater plasticity of temporal evidence representations or their attentional gain in ADHD, and greater recalibration of higher-order causal structure in TD children. ADHD is increasingly understood as involving altered interactions among association, salience, default-mode, and sensory/motor networks, especially as cognitive demands change (Shappell et al., 2021; Michael et al., 2025). This makes a network-level divergence plausible, but without neural measurements it remains a roadmap for EEG or fMRI rather than a localization claim.

The lack of cross-task correspondence suggests that the fitted parameters were task-dependent rather than trait-like. Neither *p_common_*nor ***σ****_t_* correlated across SJ and SIFI when pre- and post-training estimates were pooled. Multisensory causal inference can be feature-specific (Badde et al., 2023), and the two tasks make different demands: SJ requires an explicit judgment of simultaneity, whereas SIFI requires a numerosity report in which timing influences whether auditory evidence is allowed to alter vision. A parameter with the same mathematical label may therefore describe a task-conditioned computation rather than a context-free personal trait. The null correlations are consistent with this interpretation, although they are not decisive because parameter reliability limits cross-task correlations and the pooled analysis treats repeated observations from the same child as independent (Hedge et al., 2018; Katahira et al., 2024).

Several limitations should guide the interpretation of these findings. First, the absence of an active control condition prevents the observed pre–post changes from being attributed specifically to simultaneity-judgment training. Repeated exposure to the stimuli, task familiarization, or retesting may also have contributed to the behavioral and computational changes. The effects should therefore be described as training-related rather than definitively training-induced. Future studies should include a control condition matched for stimulus exposure, task duration, and feedback while omitting the critical temporal-discrimination contingency. Second, decision strategy was assigned according to the best-fitting model for each participant and session. The frequent transitions between model averaging, model selection, and probability matching may reflect genuine changes in decision policy, but they may also arise when competing strategies provide similar model evidence. Although no systematic group-level redistribution of strategies was observed, the stability of individual strategy classification remains uncertain. This issue could be addressed through model recovery, comparison of model weights, and hierarchical approaches that represent uncertainty over decision strategies rather than assigning each observation to a single winning model. Third, the σ*_t_* interaction is equivalent to a narrowing of an imprecisely estimated baseline difference, and in both tasks the group with higher pre-training σ*_t_* showed the larger decrease. Regression toward the mean cannot be excluded. This does not apply to *p*_common_, which was matched at baseline.

The practical implication is not simply that one group learned better. Both groups learned, but the training altered different computational targets. The broader lesson is that perceptual plasticity cannot be read directly from its behavioral endpoint. Training reduced audiovisual binding in both groups, yet the latent computation that moved was different. Whether the attenuated prior change ultimately reflects rigidity, a timescale-specific learning imbalance, altered credit assignment, or state-dependent sensory gain remains to be decided. Framing those alternatives in a shared generative model turns them from broad descriptions of ADHD into competing, testable mechanisms. Multisensory perceptual learning is therefore not a single capacity that is preserved or impaired; it is a family of adaptations, and developmental conditions can determine which member of that family carries learning.

## Data and code availability

Data and all the codes for analysis and modelling have been share via: https://github.com/evans1112/ADHD_perceptual_learning

