## Supplemental Material for "Dissociable changes in temporal precision and causal expectations following perceptual training"

**Supplementary Materials**

### **Notation and causal structure**

The latent causal variable C took two values: C = 1 indicated that the auditory and visual observations were generated by a common audiovisual source, whereas C = 2 indicated independent modality-specific sources. The prior probability of a common cause was *p_common_* = P(C = 1). Numerosity measurements were denoted *x_V_* and *x_A_* and temporal-onset measurements *t_V_* and *t_A_*. Subscripts V and A denote visual and auditory modalities, respectively.

**Supplementary Table S1.** Model notation.

| **Symbol** | **Definition** | **Unit / range** |
| --- | --- | --- |
| *S_V_, S_A_* | Physical visual and auditory numerosities | 0–2 events |
| *x_V_, x_A_* | Internal visual and auditory numerosity measurements | arbitrary numerosity units |
| *t_V_, t_A_* | Internal visual and auditory onset measurements | ms |
| *SOA* | Physical auditory-onset time relative to visual onset | ms |
| *p_common_* | Prior probability of a common audiovisual cause | 0–1 |
| *σ_V_, σ_A_* | Visual and auditory numerosity measurement noise | numerosity units |
| *σ_p_, μ_p_* | SD and mean of the numerosity prior | numerosity units |
| *σ_t_* | Shared temporal measurement noise | ms |
| *δ* | Auditory–visual latency offset added to auditory onset | ms |
| *σ_tp_, μ_t_* | SD and mean of the temporal source prior | ms |

### **Two-dimensional BCI model of the SIFI**

#### **Sensory measurements and source priors**

On each Monte Carlo trial, physical flash and beep counts generated noisy internal numerosity measurements. Visual onset defined the temporal origin. A single temporal-noise parameter was used for both modalities because separate auditory and visual temporal standard deviations are identified primarily through their quadrature sum in this design. The latency parameter *δ* allowed the temporal-evidence function to be displaced from physical synchrony.

$x_{V}\mathcal{\sim N}\left( S_{V},{\sigma_{V}}^{2} \right),\quad x_{A}\mathcal{\sim N}\left( S_{A},{\sigma_{A}}^{2} \right)$	(1)

$t_{V}\mathcal{\sim N}\left( 0,{\sigma_{t}}^{2} \right),\quad t_{A}\mathcal{\sim N}\left( SOA+\delta,{\sigma_{t}}^{2} \right)$	(2)

$s\sim\mathcal{N}\left( \mu_{p},{\sigma_{p}}^{2} \right),\quad\varphi\sim\mathcal{N}\left( \mu_{t},{\sigma_{tp}}^{2} \right)$	(3)

The numerosity prior was defined over latent event number s. The temporal prior was defined over latent event time φ. In the focal Standard 2D BCI model, *μ_t_* = 0 ms and *σ_tp_* = 300 ms were fixed; *σ_A_* = 0.33, *σ_p_* = 1.2, and *μ_p_* = 1.4 were also fixed (Zhu et al., 2024; 2026).

#### **Marginal likelihoods under common and separate causes**

For numerosity, the common-cause likelihood integrated both modality-specific measurements over a single latent source. Under separate causes, each measurement was integrated over its own independent source:

$L_{N}^{\left( 1 \right)}=p\left( x_{V},x_{A}\mid C=1 \right)=\int p\left( x_{V}\mid s \right)p\left( x_{A}\mid s \right)p\left( s \right)ds$	(4)

$L_{N}^{\left( 2 \right)}=p\left( x_{V},x_{A}\mid C=2 \right)=\left[ \int p\left( x_{V}\mid s_{V} \right)p\left( s_{V} \right)ds_{V} \right]\left[ \int p\left( x_{A}\mid s_{A} \right)p\left( s_{A} \right)ds_{A} \right]$	(5)

The temporal dimension was defined analogously:

$L_{T}^{\left( 1 \right)}=p\left( t_{V},t_{A}\mid C=1 \right)=\int p\left( t_{V}\mid\varphi\right)p\left( t_{A}\mid\varphi\right)p\left( \varphi\right)d\varphi$	(6)

$L_{T}^{\left( 2 \right)}=p\left( t_{V},t_{A}\mid C=2 \right)=\left[ \int p\left( t_{V}\mid\varphi_{V} \right)p\left( \varphi_{V} \right)d\varphi_{V} \right]\left[ \int p\left( t_{A}\mid\varphi_{A} \right)p\left( \varphi_{A} \right)d\varphi_{A} \right]$	(7)

All four integrals are Gaussian and were evaluated analytically through log likelihood ratios. For dimension d ∈ {N,T}, the dimension-specific common-cause posterior is

$q_{d}=\frac{L_{d}^{\left( 1 \right)}p_{common}}{L_{d}^{\left( 1 \right)}p_{common}+L_{d}^{\left( 2 \right)}\left( 1-p_{common} \right)}$	(8)

Following the multidimensional factorization of Zhu et al. (2026), the numerosity and temporal contributions were combined as

$w_{C}=min\left\{ 1,\frac{q_{N}q_{T}}{p_{common}} \right\}$	(9)

Equivalently, before the upper bound at one is applied,

$w_{C}=\frac{L_{N}^{\left( 1 \right)}L_{T}^{\left( 1 \right)}p_{common}}{\left[ L_{N}^{\left( 1 \right)}p_{common}+L_{N}^{\left( 2 \right)}\left( 1-p_{common} \right) \right]\left[ L_{T}^{\left( 1 \right)}p_{common}+L_{T}^{\left( 2 \right)}\left( 1-p_{common} \right) \right]}$	(10)

The implementation evaluated log(L_d_^(1)^/L_d_^(2)^) and used numerically stable logistic transforms. This avoids underflow from products of small Gaussian densities. The upper bound at one is part of the implemented multidimensional combination rule. Accordingly, w_C_ is referred to below as the common-cause weight rather than as a posterior obtained from a single joint two-hypothesis denominator.

#### **Numerosity estimates and decision rules**

Conditional on a common cause, visual and auditory measurements and the numerosity prior were combined according to their precisions. Conditional on separate causes, the task-relevant visual estimate used only the visual measurement and the prior:

$\hat{s}_{C=1}=\frac{\frac{x_{V}}{{\sigma_{V}}^{2}}+\frac{x_{A}}{{\sigma_{A}}^{2}}+\frac{\mu_{p}}{{\sigma_{p}}^{2}}}{\frac{1}{{\sigma_{V}}^{2}}+\frac{1}{{\sigma_{A}}^{2}}+\frac{1}{{\sigma_{p}}^{2}}}$	(11)

$\hat{s}_{V,C=2}=\frac{\frac{x_{V}}{{\sigma_{V}}^{2}}+\frac{\mu_{p}}{{\sigma_{p}}^{2}}}{\frac{1}{{\sigma_{V}}^{2}}+\frac{1}{{\sigma_{p}}^{2}}}$	(12)

Three mappings from causal uncertainty to the visual estimate were considered. Model averaging used *ŝ_V_* = *w_C_ ŝ_C=1_* + (1−*w_C_*)*ŝ_V,C=2_*. Model selection used the common-cause estimate when *w_C_* > 0.5 and the separate-cause estimate otherwise. Probability matching sampled the common-cause estimate with probability *w_C_*. The continuous estimate was mapped to the binary task response by reporting two flashes when *ŝ_V_* ≥ 1.5 and one flash otherwise. The predicted probability of a two-flash response was the proportion of Monte Carlo trials crossing this threshold. To keep binomial log likelihoods finite, predicted probabilities were transformed as *p** = 0.0005 + 0.999*p*, equivalent to a fixed 0.1% symmetric lapse mixture.

For unisensory conditions, the likelihood standard deviation of the absent modality was set to 1,000, rendering that measurement uninformative. Measurement noise was nevertheless generated using the model parameter before this likelihood override, preserving the visual-only trials that directly constrain *σ_V_*.

### **Temporal BCI model of simultaneity judgments**

The SJ model retained only the temporal dimension. The same temporal measurement model, temporal source prior, *p_common_*, *σ_t_*, and *δ* definitions were used, but parameters were fitted independently of the SIFI model. The common-cause posterior was

$P\left( C=1\mid t_{V},t_{A} \right)=\frac{L_{T}^{\left( 1 \right)}p_{common}}{L_{T}^{\left( 1 \right)}p_{common}+L_{T}^{\left( 2 \right)}\left( 1-p_{common} \right)}$	(13)

With equal temporal noise, define *d* = *t_A_* − *t_V_* and M = (*t_V_* + *t_A_*)/2 − *μ_t_*. The temporal log evidence ratio reduces to

$R_{T}=log\left[ \frac{L_{T}^{\left( 1 \right)}}{L_{T}^{\left( 2 \right)}} \right]=K+Ad^{2}+BM^{2}$	(14)

$A=\frac{-1}{4{\sigma_{t}}^{2}}+\frac{1}{4\left( {\sigma_{t}}^{2}+{\sigma_{tp}}^{2} \right)},\quad B=\frac{1}{\left( {\sigma_{t}}^{2}+{\sigma_{tp}}^{2} \right)}-\frac{1}{\left( {\sigma_{t}}^{2}+2{\sigma_{tp}}^{2} \right)}$	(15)

$K=log\left\{ \frac{{\sigma_{t}}^{2}+{\sigma_{tp}}^{2}}{\sigma_{t}\sqrt{{\sigma_{t}}^{2}+2{\sigma_{tp}}^{2}}} \right\}$	(16)

The posterior was logistic[*R_T_* + *logit*(*p_common_*)]. Under model selection, a simultaneous response occurred when this posterior exceeded 0.5; its response probability was evaluated by one-dimensional quadrature over M. Under probability matching, the response probability was the expected posterior and was evaluated by two-dimensional quadrature over d and M. The final prediction included a fixed lapse mixture, *p_obs_* = 0.02(0.5) + 0.98*p_model_*. Thus, lapse trials comprised 2% of responses and were unbiased between “simultaneous” and “asynchronous.”

### **Candidate SIFI architectures and parameter constraints**

The focal Standard 2D BCI model was compared with four targeted alternatives. Internal implementation labels are omitted here; the scientific names below should be used in the manuscript and supplement.

**Supplementary Table S2.** Candidate BCI model architectures.

| **Architecture** | **Free parameters** | **Fixed / externally supplied values** | **Purpose** |
| --- | --- | --- | --- |
| Standard 2D BCI (focal) | *p_common_*, *σ_V_*, *σ_t_, δ* | *σ_A_* = 0.33, *σ_p_* = 1.2, *μ_p_* = 1.4 | Primary decomposition of causal prior, visual noise, temporal noise, and latency. |
| Fully flexible sensory–temporal | *p_common_, σ_V_, σ_A_, σ_t_, δ* | *σ_p_* = 1.2, *μ_p_* = 1.4 | Tests whether freeing auditory uncertainty and both temporal parameters improves fit. |
| Cross-task latency-constrained | *p_common_, σ_V_, σ_A_, σ_t_* | *δ* fixed to the participant’s SJ estimate; *σ_p_* = 1.2, *μ_p_* = 1.4 | Tests free SIFI temporal noise while constraining latency across tasks. |
| Fixed-temporal flexible-auditory | *p_common_, σ_V_, σ_A_, σ_p_, μ_p_* | *σ_t_* = 60 ms; *δ* = 0 | Tests whether numerosity flexibility can replace individual temporal parameters. |
| Fixed-temporal constrained-auditory | *p_common_, σ_V_, σ_p_, μ_p_* | *σ_t_* = 60 ms; *δ* = 0; *σ_A_* = 0.33 | Tests whether flexible temporal parameters are required when auditory uncertainty is fixed. |

### **Likelihood, optimization, and comparison**

Models were fitted separately for every participant and pre/post session. Missed or invalid responses were excluded before aggregation. For condition *j, k_j_* denotes the number of target responses among n_j_ valid trials and *p_j_(θ)* is the model prediction. Parameters minimized the binomial negative log likelihood

$NLL\left( \theta\right)=-\sum_{j} \left[ k_{j}logp_{j}\left( \theta\right)+\left( n_{j}-k_{j} \right)log\left( 1-p_{j}\left( \theta\right) \right) \right]$	(17)

SIFI predictions used 1,000 Monte Carlo samples per condition. Standard-normal and probability-matching draws were generated once and reused at every objective evaluation (common random numbers), making the sampled objective deterministic. SJ predictions were evaluated by deterministic quadrature. Bayesian Adaptive Direct Search (BADS; Acerbi and Ma, 2017) used five randomized starts per decision rule. The lowest-NLL solution was retained. Within an architecture, the winning decision rule minimized

$BIC=klogN+2NLL$	(18)

where k is the number of free parameters and *N* is the number of valid trials. Because decision rules within an architecture shared *k*, this comparison was equivalent to selecting the lowest NLL. Architecture comparisons were restricted to the 100 participant-by-session cells with all candidate fits. Paired model differences were evaluated by a Friedman test followed by paired Wilcoxon signed-rank tests with Holm correction.

For SIFI, two-fold cross-validation alternated trials within each stimulus condition so that both folds contained every condition. The decision rule selected from the complete dataset was retained, parameters were re-estimated on one fold using three restarts, and held-out binomial log likelihood was accumulated over the other fold. This procedure was used as an out-of-sample diagnostic rather than as the primary architecture-selection criterion.

### **Goodness-of-fit and identifiability diagnostics**

Raw R² was the squared Pearson correlation between observed and predicted condition-level response proportions. Because observed proportions contain binomial sampling noise, an estimated noise ceiling was also computed:

${R^{2}}_{ceiling}=\frac{max\left\{ Var\left( p_{obs} \right)-mean\left[ \frac{p_{obs}\left( 1-p_{obs} \right)}{n} \right],0 \right\}}{Var\left( p_{obs} \right)}$	(19)

When *R²_ceiling_* > 0.15, *R²_normalized_* = R²/R²_ceiling_ was reported. Model deviance relative to the saturated binomial model was

$G^{2}=2\left( {NLL}_{model}-{NLL}_{saturated} \right),\quad df=J-k$	(20)

A non-significant deviance test indicates that remaining discrepancies are compatible with binomial sampling error. We also counted solutions within 1% of the width of a hard bound (*p_common_* ≤ .011 or ≥ .989; *σ_V_* ≤ .051 or ≥ 2.99; *σ_t_* ≤ 5.1 or ≥ 490 ms; |*δ*| ≥ 294 ms) and recorded how many randomized starts reached an NLL within 0.01 of the best solution.

### **Parameter recovery**

Recovery was conducted for the focal Standard 2D BCI model under model averaging, the decision rule used for the primary parameter analysis. One hundred synthetic datasets were generated using the same 24 SIFI conditions as the empirical design, with 20 trials per condition. Generating probabilities were evaluated with 10,000 Monte Carlo samples per condition, after which binomial response counts were sampled. Each dataset was then fitted with the same model-averaging rule using 1,000 Monte Carlo samples per condition, common random numbers, and five randomized BADS starts. Recovery was summarized using Pearson correlations, Spearman rank correlations with 10,000-sample percentile-bootstrap 95% confidence intervals, bias, mean absolute error (MAE), and root-mean-square error (RMSE). We additionally compared the generating and recovered condition-probability surfaces. Because all datasets were generated and fitted under model averaging, this analysis evaluated recovery of the four focal parameters rather than recovery of decision strategies.

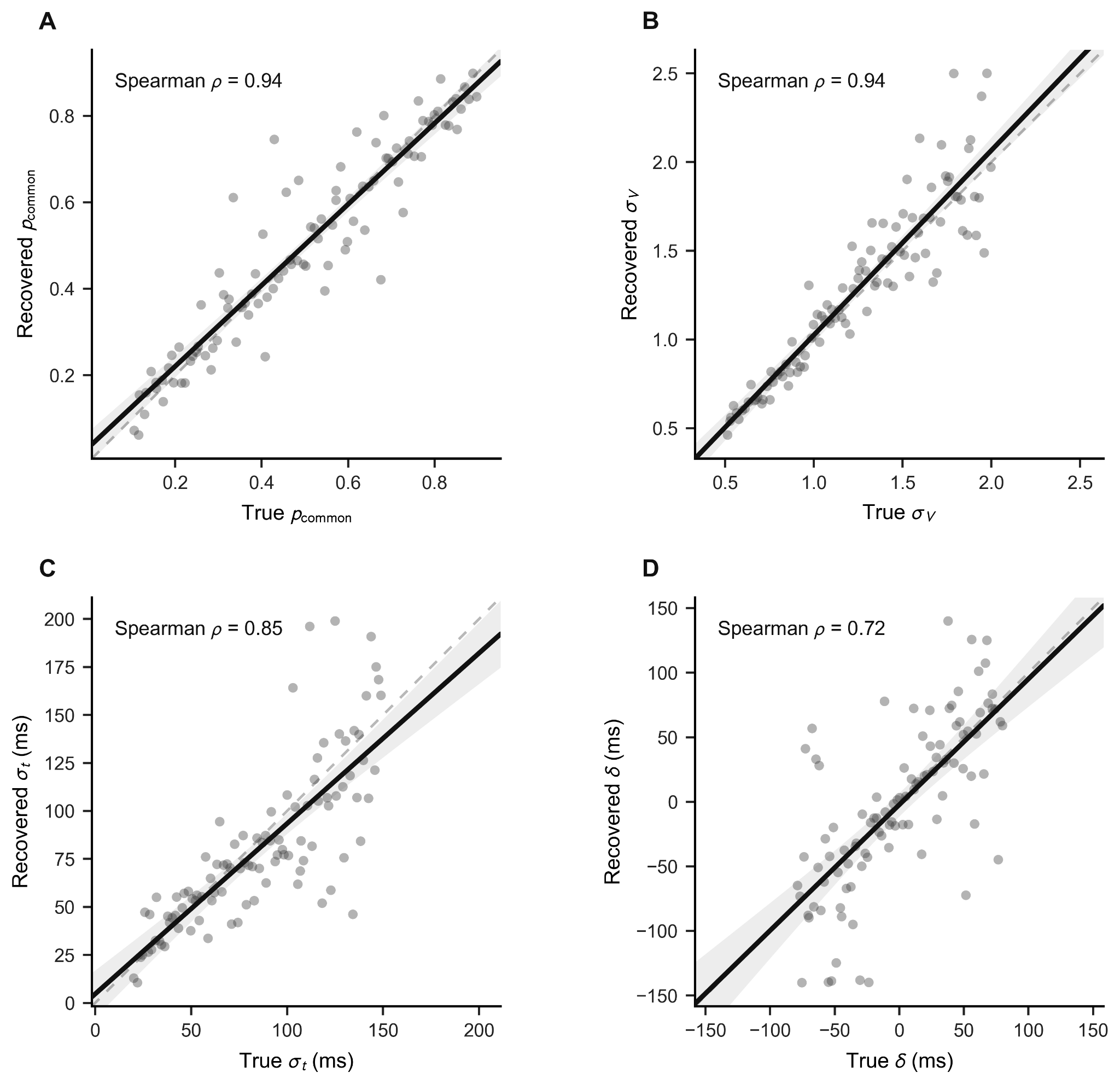

**Figure S1. Parameter recovery for the Standard 2D Bayesian causal-inference model.** Recovery was evaluated using 100 synthetic datasets generated and fitted under model averaging, with 20 trials per stimulus condition. Each point represents one simulated dataset. Recovered estimates are plotted against the generating values for **(A)** the common-cause prior, *p_common_*; **(B)** visual numerosity uncertainty, *σ_V_*; **(C)** temporal uncertainty, *σ_t_*; and **(D)** the auditory–visual latency offset, *δ*. Solid black lines show linear regression fits, shaded regions indicate 95% confidence intervals, and dashed grey lines denote perfect recovery (identity). Spearman rank correlations are reported within each panel. Recovery was strongest for *p_common_* and *σ_V_*, and remained strong for *σ_t_*, and δ.

**Supplementary Table S3. Prediction-level recovery diagnostics.**

| **Diagnostic** | **Value** |
| --- | --- |
| Median probability-surface R² | .987 |
| Median probability-surface RMSE | .028 |

All synthetic datasets were generated and fitted using model averaging; decision-strategy recovery was not evaluated.

Parameter recovery was strong across the four focal parameters. Spearman correlations were .944 for *p_common_* (95% bootstrap CI [.899, .968]), .944 for *σ_V_* ([.900, .966]), .846 for *σ_t_* ([.754, .907]), and .723 for *δ* ([.578, .835]). Bias was small for *p_common_* (.002), *σ_V_* (.036), *σ_t_* (−4.87 ms), and *δ* (−2.21 ms). Absolute recovery error was lowest for *p_common_* (MAE = .052) and *σ_V_* (MAE = .125); temporal-noise and latency estimates were less precise in their native units but remained identifiable (*σ_t_*: MAE = 17.64 ms; *δ*: MAE = 30.44 ms). Recovered parameters closely reproduced the generating response surfaces (median probability-surface *R²* = .987; median *RMSE* = .028). Thus, the implemented design supports recovery of the group-level parameter structure used in the main analysis.

### **Statistical analysis of fitted parameters**

For the SIFI model, the common-cause prior *p_common_* and temporal uncertainty *σ_t_* were designated as the two theoretically focal computational outcomes. Group differences in post-minus-pre change scores were evaluated using two-sided Welch tests, with Holm adjustment across the two focal Group × Time tests. Because both interactions survived correction, the four corresponding within-group simple effects, two parameters in each of the two groups, were evaluated using two-sided Wilcoxon signed-rank tests and Holm-adjusted as a separate family. The visual-to-auditory sensory-noise ratio was treated as a secondary outcome; its two within-group tests were Holm-adjusted together, whereas its single Group × Time test formed a separate one-test family. For the secondary SJ analyses, Holm adjustment was applied across the six group-by-parameter within-group tests and separately across the three Group × Time tests. Confidence intervals for mean changes and group differences were obtained using the corresponding paired or Welch *t* framework. The SIFI latency offset *δ* was treated as a nuisance parameter and was not included in the focal correction families.

Cross-task correlations followed the requested descriptive analysis: pre- and post-training values were pooled, giving 100 participant-session observations, and Pearson correlations were calculated between SIFI and SJ estimates of *p_common_* and *σ_t_*. Because the same child contributed two observations, these p values do not constitute cluster-robust inference. Decision strategies were assigned by the lowest BIC within the focal SIFI architecture; pre/post marginal homogeneity was assessed separately by group using Stuart–Maxwell tests.

**Statistical reporting and reproducibility**

All behavioural analyses were implemented in Python using NumPy, pandas, SciPy, and custom scripts. Unless stated otherwise, tests were two-sided (*α* = 0.05), estimates are accompanied by 95% confidence intervals, and Holm correction controlled the families defined above. No observations were removed based on outcome magnitude or direction.

### **Complete focal-model parameter results**

**Supplementary Table S4.** Complete focal-model parameter results. Pre/post changes in computational parameters reported in the main analysis.

| **Task** | **Parameter** | **Effect** | **Group** | **Pre mean** | **Post mean** | **Estimate** | **Adj. p** |
| --- | --- | --- | --- | --- | --- | --- | --- |
| SJ | *p*_common_ | Post − pre | TD | 0.585 | 0.399 | −0.186 | .053 |
| SJ | *p*_common_ | Post − pre | ADHD | 0.633 | 0.549 | −0.083 | .117 |
| SJ | *p*_common_ | Group × time | ADHD − TD | — | — | 0.103 | .398 |
| SJ | *σ*_t_ | Post − pre | TD | 188.556 | 129.648 | −58.908 | .679 |
| SJ | *σ*_t_ | Post − pre | ADHD | 165.183 | 180.404 | 15.221 | .808 |
| SJ | *σ*_t_ | Group × time | ADHD − TD | — | — | 74.129 | .310 |
| SJ | *δ* | Post − pre | TD | 53.127 | 25.587 | −27.540 | .808 |
| SJ | *δ* | Post − pre | ADHD | 79.839 | 103.324 | 23.485 | .808 |
| SJ | *δ* | Group × time | ADHD − TD | — | — | 51.025 | .398 |
| SIFI | *p*_common_ | Post − pre | TD | 0.637 | 0.379 | −0.259 | **.021*** |
| SIFI | *p*_common_ | Post − pre | ADHD | 0.631 | 0.630 | −0.001 | 1.000 |
| SIFI | *p*_common_ | Group × time | ADHD − TD | — | — | 0.258 | **.048*** |
| SIFI | *σ*_t_ | Post − pre | TD | 114.097 | 115.491 | 1.393 | 1.000 |
| SIFI | *σ*_t_ | Post − pre | ADHD | 228.862 | 110.194 | −118.668 | **.006*** |
| SIFI | *σ*_t_ | Group × time | ADHD − TD | — | — | −120.062 | **.048*** |
| SIFI | *σ*_V_*/σ*_A_ | Post − pre | TD | 4.547 | 4.977 | 0.430 | 1.000 |
| SIFI | *σ*_V_*/σ*_A_ | Post − pre | ADHD | 4.449 | 4.268 | −0.181 | 1.000 |
| SIFI | *σ*_V_*/σ*_A_ | Group × time | ADHD − TD | — | — | −0.611 | .532 |

**Supplementary Table S5.** Baseline group comparisons of focal-model parameters before training.

| **Task** | **Parameter** | **TD M (SD)** | **ADHD M (SD)** | **Welch t (df)** | **p** |
| --- | --- | --- | --- | --- | --- |
| SIFI | *p*_common_ | 0.637 (0.309) | 0.631 (0.265) | −0.08 (47.99) | .940 |
| SIFI | *σ*_V_ | 1.500 (0.868) | 1.468 (0.830) | −0.13 (47.32) | .894 |
| SIFI | *σ*_t_ | 114.097 (140.112) | 228.862 (170.582) | 2.57 (42.64) | **.014*** |
| SIFI | *δ* | 28.609 (120.826) | 88.454 (176.219) | 1.38 (37.99) | .177 |
| SJ | *p*_common_ | 0.585 (0.245) | 0.633 (0.180) | 0.79 (47.09) | .434 |
| SJ | *σ*_t_ | 188.556 (151.881) | 165.183 (123.763) | −0.60 (47.92) | .552 |
| SJ | *δ* | 53.127 (152.362) | 79.839 (117.456) | 0.70 (47.57) | .488 |

**TOJ pre-post report**

The TOJ psychometric functions showed evidence of a post-training response-criterion shift. In TD children, the proportion of visual-first responses increased from 0.48 to 0.70, mean change = 0.21, 95% CI [0.10, 0.33], *t*(26) = 3.97, *p* < .001, *d*_z_ = 0.76, adjusted *p* = .002. The corresponding increase in children with ADHD, from 0.53 to 0.60, was smaller and did not reach significance, mean change = 0.07, 95% CI [−0.02, 0.17], *t*(22) = 1.55, *p* = .136, *d*_z_ = 0.32. The nominal Group × Time interaction, *t*(47.8) = −2.00, *p*= .051, did not survive Holm correction, adjusted *p* = .103. Because this response shift was not accompanied by improved accuracy or a narrower temporal window, it is more consistent with a change in response criterion than with enhanced temporal-order discrimination.

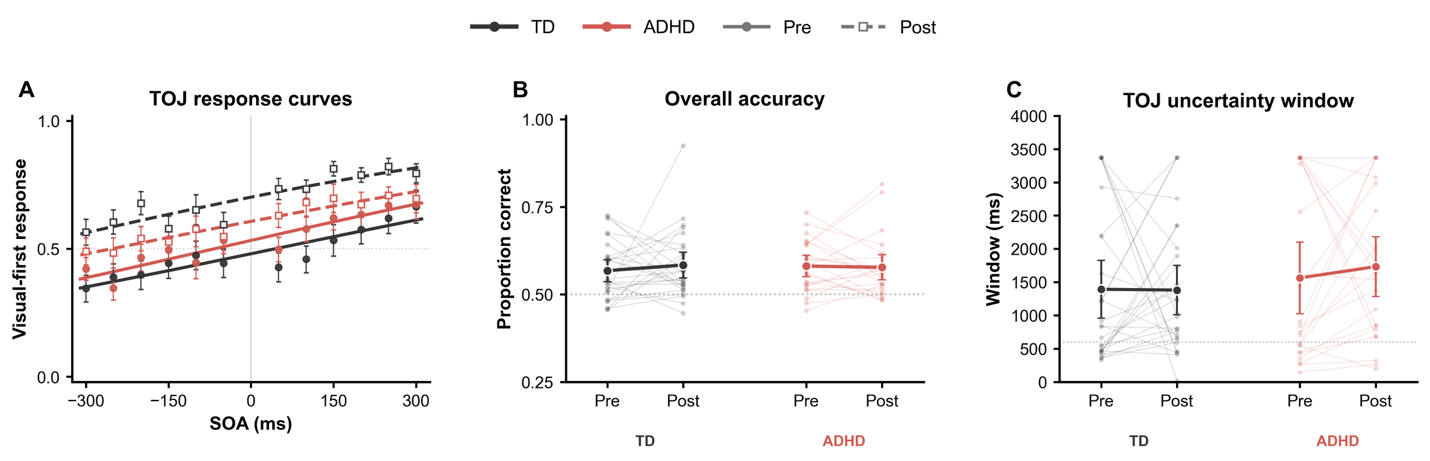

**Figure S2. Temporal-order-judgment performance before and after training.**
(A) The proportion of visual-first responses as a function of stimulus-onset asynchrony (SOA). Symbols show group means ± SEM, and smooth lines show cumulative Gaussian functions fitted to the participant-averaged data. Negative SOAs indicate auditory-leading trials and positive SOAs indicate visual-leading trials. The horizontal dotted line marks an equal probability of visual-first and auditory-first responses, and the vertical line indicates physical synchrony. (B) Overall TOJ accuracy for each participant before and after training. The horizontal dotted line indicates chance performance. (C) Individual TOJ uncertainty windows, defined as the interval between the 25% and 75% points of each participant’s fitted cumulative Gaussian function. The horizontal dotted line indicates the 600-ms tested SOA span; estimates above this value require extrapolation beyond the observed stimulus range and should therefore be interpreted cautiously. In B and C, small points and thin lines represent individual participants, large points represent group means, and error bars indicate 95% confidence intervals. TD participants are shown in black and participants with ADHD in red; pre-training measurements are represented by solid lines and filled circles, and post-training measurements by dashed lines and open squares.
